# Rab11 is essential to pancreas morphogenesis, lumen formation and endocrine mass

**DOI:** 10.1101/2022.11.17.516976

**Authors:** Haley R. Barlow, Yadanar Htike, Luke Fassetta, Neha Ahuja, Tyler Bierschenk, D. Berfin Azizoglu, Juan Flores, Nan Gao, Denise Marciano, Ondine Cleaver

## Abstract

The molecular links between tissue-level morphogenesis and the differentiation of cell lineages in the pancreas remain elusive despite a decade of studies. We previously showed that in pancreas both these processes depend on proper lumenogenesis. The Rab GTPase Rab11 has been shown to be essential to epithelial lumen formation *in vitro*, however few studies have addressed its functions *in vivo* and none have tested its requirement in pancreas. Here, we show that Rab11 is critical to proper pancreas development. Co-deletion of the Rab11 isoforms *Rab11A* and *Rab11B* in the developing pancreatic epithelium (Rab11^pancDKO^) results in ~50% neonatal lethality, and surviving adult Rab11^pancDKO^ mice exhibit defective endocrine function. Loss of Rab11 in the embryonic pancreas results in morphogenetic defects of the epithelium linked to defective lumen formation and interconnection. In contrast to wildtype cells, Rab11^pancDKO^ cells attempt to form multiple lumens, resulting in a failure to coordinate a single apical membrane initiation site (AMIS) between groups of cells. We show that these defects are due to failures in vesicle trafficking, as apical components remain trapped within Rab11^pancDKO^ cells. Together, these observations suggest Rab11 directly regulates epithelial lumen formation and morphogenesis. Our report links intracellular trafficking to organ morphogenesis *in vivo*, and presents a novel framework for decoding pancreatic development.

**HIGHLIGHTS:**

- Rab11A^f/f^;Rab11B^-/-^;Pdx1-Cre pancreas displays disruption of epithelial organization and reduction of endocrine cell mass.
- Loss of Rab11 results in disruption of pancreatic lumen continuity due to a failure of lumen formation.
- Epithelial cells lacking Rab11 display abnormal polarity.

## INTRODUCTION

Lumen formation is a critical process in multicellular organism development, as tubular structures allow for the distribution of gases and liquids deep into organs and tissues. Indeed, the formation of a lumen within a tubular structure is the functional rate-limiting step in the formation of many organ systems including the kidney, lung and cardiovascular systems. Somewhat surprisingly, different tissues utilize different cellular strategies to form lumens (Andrew and Ewald, 2010; Marciano, 2017). Lumens can form by cavitation (e.g. pro-amniotic lumen) (Meng et al., 2017); entrapment (e.g. *Drosophila* heart) (Rugendorff et al., 1994); wrapping (e.g. neural tube formation) (Harland, 1994); cell hollowing (e.g. Drosophila trachea) (Samakovlis et al., 1996); or cord hollowing, in which multiple cells coordinate to target apical proteins to a central location and form a lumen (e.g. initial aortic lumen) (Strilic et al., 2009). Probing the molecular mechanisms driving tissue lumen formation *in vivo* will likely be critical to therapeutic efforts to grow human tissue *ex vivo* for replacement therapies, since the presence and proper organization of lumens is essential to the function of organs (Ryan and Cleaver, 2022).

The cord-hollowing, or *de novo*, mechanism of lumen formation has been observed in organ systems including the cardiovascular system, the kidney, and the pancreas (Gao et al., 2017; Strilic et al., 2009; Villasenor et al., 2010). However, the molecular machinery underlying *de novo* lumen formation has been most thoroughly examined *in vitro* using Madin Darby Canine Kidney (MDCK) cells (Bryant et al., 2010). In this system, a new single central lumen was shown to be initiated between cells following the coordination of a single <u>a</u>pical <u>m</u>embrane <u>i</u>nitiation <u>s</u>ite (termed AMIS) (Bryant et al., 2010). AMIS formation is a dynamic process, as membrane-associated proteins are rapidly moved to and from various locations as cells acquire polarity. Apically and basally localized proteins are trafficked, junctions are rearranged, cell shape is changed, and the cytoskeleton undergoes rapid and dynamic changes to support these processes. Moreover, components of polarity determining complexes such as the Par (apical), Crumbs (apical) and Scribble (basolateral) complexes, which localize to the apical or basolateral membranes as polarity is established, are known to contribute to AMIS formation (Overeem et al., 2015). These complexes regulate strong positive feedback loops that maintain the distinct identities of the apical and basolateral membranes (for further review see Roman-Fernandez and Bryant, 2016). Although much is known about AMIS and lumen formation in cultured cells, the molecular machinery that drives lumen formation in developing tissues remains largely unexplored.

The developing murine pancreas presents an opportunity to study *de novo* lumen formation *in vivo*. The pancreatic epithelium first buds from the foregut endoderm as a stratified epithelial placode around embryonic day (E) 8.75 (Flasse et al., 2021; Pictet et al., 1972; Slack, 1995; Villasenor et al., 2010). Within this multilayered epithelium, *de novo* microlumens emerge in the initial, unbranched pancreas (Braitsch et al., 2019; Villasenor et al., 2010). These microlumens go on to connect to each other and to the foregut lumen, forming a highly interconnected lumenal plexus, as the pancreas grows and ramifies into a branching, glandular epithelial tree (Hick et al., 2009; Kesavan et al., 2009; Villasenor et al., 2010). Unlike other organ systems, *de novo* lumenogenesis continues to occur alongside lumen remodeling and extension as the pancreas grows and ramifies into a branching, glandular epithelial tree.

How are these dynamic cellular activities carried out and coordinated? We chose to focus on how vesicle-mediated recycling regulates membrane and protein trafficking during lumen formation and pancreas morphogenesis. Specifically, we honed in on the master of membrane recycling Rab11 (Rab11A and Rab11B). Rab11 is a member of the Ras superfamily of GTPases that is known to regulate secretory and endocytic vesicular trafficking, including recycling of a number of membrane surface and polarity proteins (Goldenring et al., 1996; Ossipova et al., 2014; Zhang et al., 2021). Rab11 has been shown to regulate cell polarity and cell division in yeast and zebrafish (Tanasic et al., 2022) as well as lumen formation in MDCK cells (Desclozeaux et al., 2008; Hales et al., 2002; Neto et al., 2013; Overeem et al., 2015; Rathbun et al., 2020; Roland et al., 2011). During MDCK lumen formation, Rab11 localizes at the AMIS, and traffics proteins from the basal to the apical membrane as cells establish polarity. In the absence of Rab11, lumen formation is disrupted (Bryant et al., 2010; Desclozeaux et al., 2008). These observations established Rab11 as an important regulator of polarity determination and lumen formation in cultured epithelial cells. However, there has been surprisingly little examination of the role of Rab11 during mammalian *in vivo* lumen formation. Our studies have previously implicated Rab11 in pancreatic lumen formation *in vivo* (Azizoglu et al., 2017). In the absence of the junctional/scaffolding protein Afadin, the formation and organization of lumens in the pancreas is severely disrupted. Loss of Afadin leads to the disruption of Rab11 localization and to a concomitant accumulation of intracellular cargo containing both apical and junctional components. Based on these observations, we asked how Rab11 regulates pancreatic lumen formation in vivo.

This study is the first to characterize ablation of Rab11 (Rab11A and Rab11B) *in vivo* during organogenesis, and identifies functional links between vesicle trafficking, polarity acquisition, lumen formation, and tissue morphogenesis. Here we show that loss of Rab11 in the mouse pancreatic epithelium leads to postnatal lethality, failure to thrive and defective pancreatic endocrine function. These deficiencies are linked to severe fetal pancreatic hypoplasia and to a decrease in endocrine cell mass. Furthermore, the loss of Rab11 leads to profound epithelial morphogenetic defects. Nascent lumens fail to connect and form a plexus in the Rab11^pancDKO^ at E14.5, as apical proteins remain trapped inside epithelial cells. Rab11^pancDKO^ cells also display severely disrupted polarity and often participate in more than one AMIS between neighbors. This is associated with mislocalization of Par and Crumbs polarity complex proteins, as well as tight junction components like ZO-1. We propose that Rab11 is required for the formation of a single AMIS that matures into a single coordinated lumen in the pancreatic epithelium *in vivo*. When these processes are disrupted, the morphogenesis of the pancreas as a whole is defective, and endocrine cell fate is affected. Together, these findings identify cellular and molecular mechanisms underlying pancreatic morphogenesis and underscore the functional connections between lumen formation and epithelial morphogenesis.

## RESULTS

### Rab11 is expressed in the developing pancreas

We previously showed that the junctional protein Afadin was required for both lumen formation and normal localization of Rab GTPases, as well as subsequent proper pancreas development (Azizoglu et al., 2017). In this study, we assess whether Rab11 is involved in pancreas development.

We first assessed expression of Rab11A throughout pancreatic development. We primarily found it expressed in the pancreatic epithelium, with low to undetectable expression in the pancreatic mesenchyme. At all stages examined (embryonic day (E) 10.5, E11.5, E12.5, E14.5, E18.5), Rab11A was enriched at the apical membrane of nascent and established lumens (**Fig. 1 A-D’, F-H**). Throughout the progression of developmental time, there was an increase in signal intensity and coordination of localization towards the apical membrane as lumens matured (**Fig. 1 E**). We also evaluated expression in the pancreatic lineages – acinar, ductal and endocrine. Before birth at E18.5, we found Rab11A largely restricted to the apical membranes of both acinar and ductal cells (**Fig. 1 F-G’; Fig. S1 A-B’, D-D’**), while exhibiting low expression in endocrine cells (**Fig. 1 H-H’; Fig. S1 C-D’**). We confirmed the localization of Rab11A in the three main pancreatic lineages at E18.5 both by epithelial morphology (**Fig. 1 F-H’; Fig. S1 D-D’**) and by analysis of lineage-specific markers CPA1 (acinar; **Fig. S1 A-A’**), Sox9 (ductal; **Fig. S1 B-B’**), and Insulin and Glucagon (endocrine; **Fig. S1 C-C’**). Alternative immunostaining methods utilized to enhance nuclear antigen signal also revealed strong lateral localization of Rab11A in ductal cells (**Fig. S1 B-B’**), but this was not observed using any other staining method. Overall, these results support the idea that Rab11A is a strong candidate regulator of pancreatic lumen formation.

**Figure 1.**
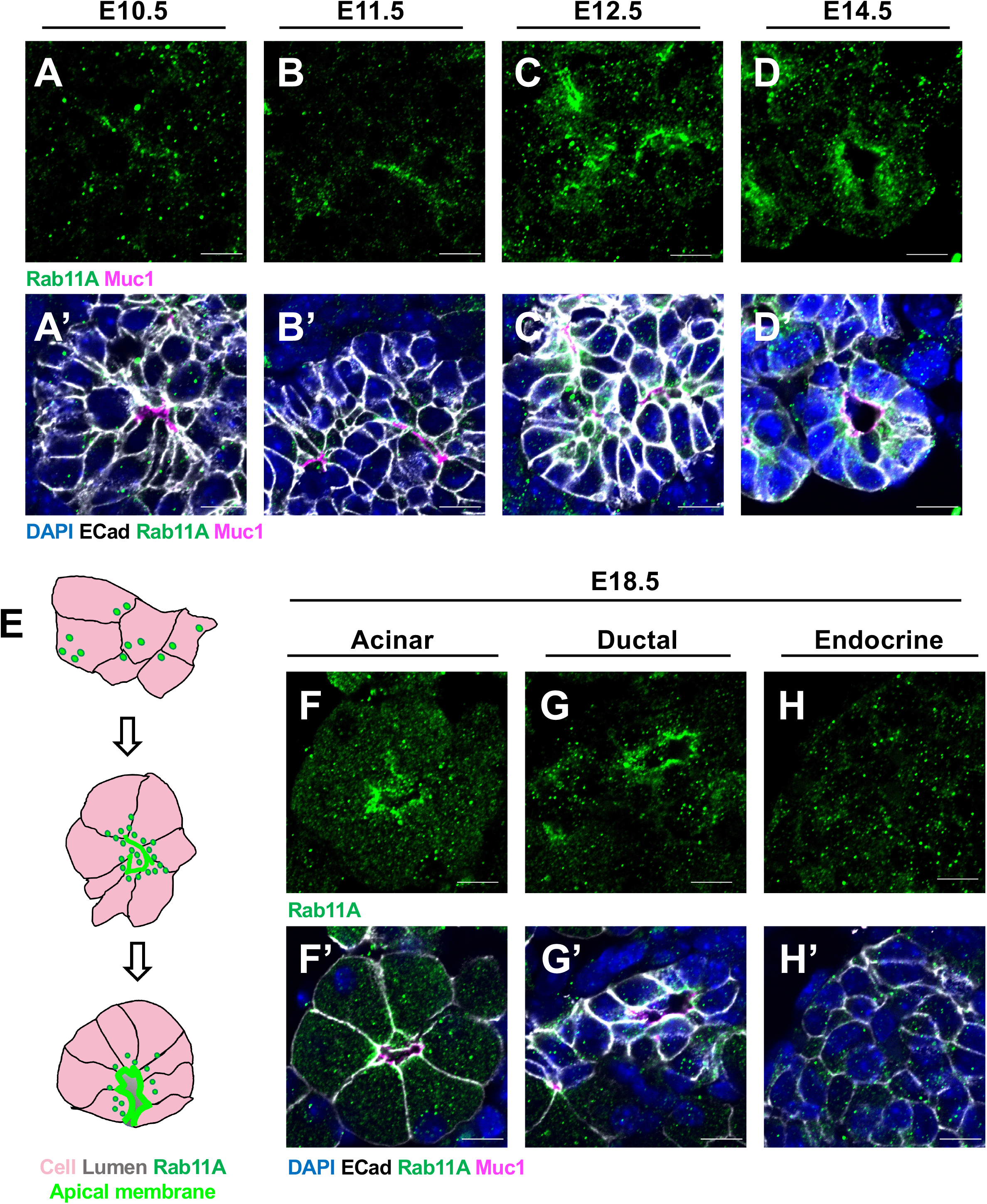
Vesicle-mediated recycling regulator Rab11 localizes to lumens throughout pancreatic development. Rab11A (green) is enriched at apical membranes (Muc1, magenta) at early- (A-C’) and mid-gestational (D-D’) stages of pancreatic development. The shift in localization of Rab11A as *de novo* lumens form is represented in (E), in which epithelial cells are pink, Rab11A^+^ vesicles are dark green, apical membrane is bright green, and the open lumen is grey. At E18.5, Rab11A localization remains restricted to apical membranes of lumens in acinar cells (F-F’) and ductal cells (G-G’). Endocrine cells (H-H’) also display low levels of Rab11 expression. n>3, scale bars = 10uM.

### Loss of Rab11 from the pancreas results in postnatal lethality

To assess a potential role of Rab11 during pancreas development, we genetically ablated *Rab11A* from the pancreatic epithelium utilizing the *Pdx1-Cre* driver in a *Rab11B*-null background (Rab11^pancDKO^) (Hingorani et al., 2003; Yu et al., 2014). *Rab11A^f/f^; Rab11B*^-/-^ females were crossed to males that were heterozygous either for *Rab11A* or *Rab11B (Rab11A^f/+^;Rab11B*^-/-^ or *Rab11A^f/f^;Rab11B^+/-^*) and had a copy of *Pdx1-Cre*. Individual mutants of *Rab11A* or *Rab11B* did not display any noticeable phenotype and were therefore included in the pool of controls for experiments, and the breeding heterozygous males indicated above were viable and fertile with no obvious abnormalities.

Expected ratios of mutants were found during embryonic development. However when we allowed pregnant dams to give birth we found that only approximately one half of the mutant animals survived, with the most common survival time being only to postnatal day 2 (P2) (**Fig. 2 A**). At postnatal day (P) 8, surviving Rab11^pancDKO^ pups exhibited clear defects in overall growth (**Fig. 2 B**). At weaning (P28), there was significant variability in animal weight and size, suggesting that the mutant animals that survive may have a less severe phenotype than those that die (**Fig. 2 C**). When subjected to a glucose tolerance test, adult Rab11^pancDKO^ males and females took longer to return to normal blood glucose levels, indicating that there are problems with endocrine function in the Rab11^pancDKO^ mice (**Fig. 1 D**). Interestingly, analysis of the variability in blood glucose measurements revealed a dramatic increase in variability in the mutant mice, again suggesting variability in phenotype severity (**Fig. 1 E**). We confirmed that *Rab11* was efficiently ablated in our mutant mice by testing the presence of Rab11A protein via immunostaining in the midgestational E14.5 pancreatic epithelium (**Fig. S2 A-B”**).

**Figure 2.**
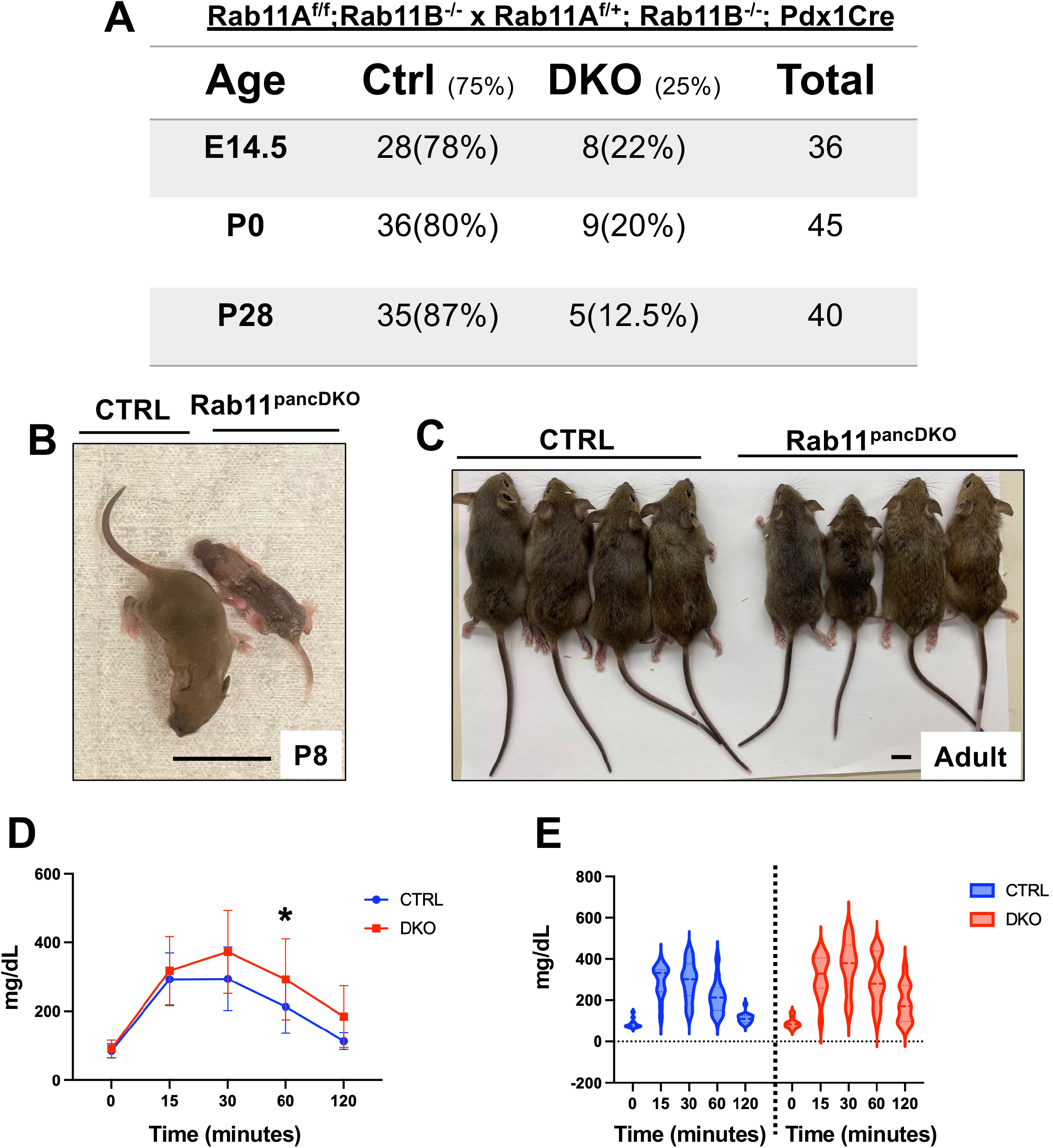
Loss of Rab11 leads to postnatal lethality, failure to thrive and defects in endocrine function. Rab11^pancDKO^ mutants are present during gestation and at birth at expected ratios (A). However, approximately half of Rab11^pancDKO^ mutants die before weaning, with many of them dying by postnatal day (P) 2 (data not shown, A). Mutants that survive beyond P2 are smaller than their littermates and display a failure to thrive (B). Of the mutants that survive to weaning (P28), Rab11^pancDKO^ mice are smaller with significantly decreased body weight at weaning (C-D). There is notable variability in phenotype severity of the adult Rab11^pancDKO^ mice (C). Despite this variability, the glucose tolerance of Rab11^pancDKO^ mice is significantly affected (E). n=111 mice analyzed for genotypes; n>3 mutants that fail to thrive postnatally; n=8 female mice and at least n=8 male mice per group for P28 analyses; error bars = standard error of the mean for (D), standard deviation for (E); scale bars = 1cm; data in (D) analyzed in PRISM by Student’s t-test; data in (E) analyzed in PRISM by 2-Way ANOVA.

### Rab11^pancDKO^ exhibit pancreas hypoplasia and epithelial defects

To assess the underlying cause of low survival rate and abnormal endocrine function in Rab11 mutant mice, we examined the pancreata of E18.5 and P2 mutants. Upon dissection of the midgut organs such as the stomach, duodenum and pancreas, we observed clear developmental defects in Rab11^pancDKO^ pancreata (**Fig. 3 A-B; Fig S3 F-I’**), accompanied by a statistically significant decrease in pancreas weight (**Fig. 3 C**). In addition to a smaller epithelial mass, the pancreas was less branched and some large cysts in the pancreatic epithelium were observed (**Fig. S3 G’, I’**, yellow arrowheads). This hypoplasia seems was specific to the pancreas, as E18.5 Rab11^pancDKO^ mutant embryos were the same size as their control littermates (**Fig. S3 E**).

**Figure 3.**
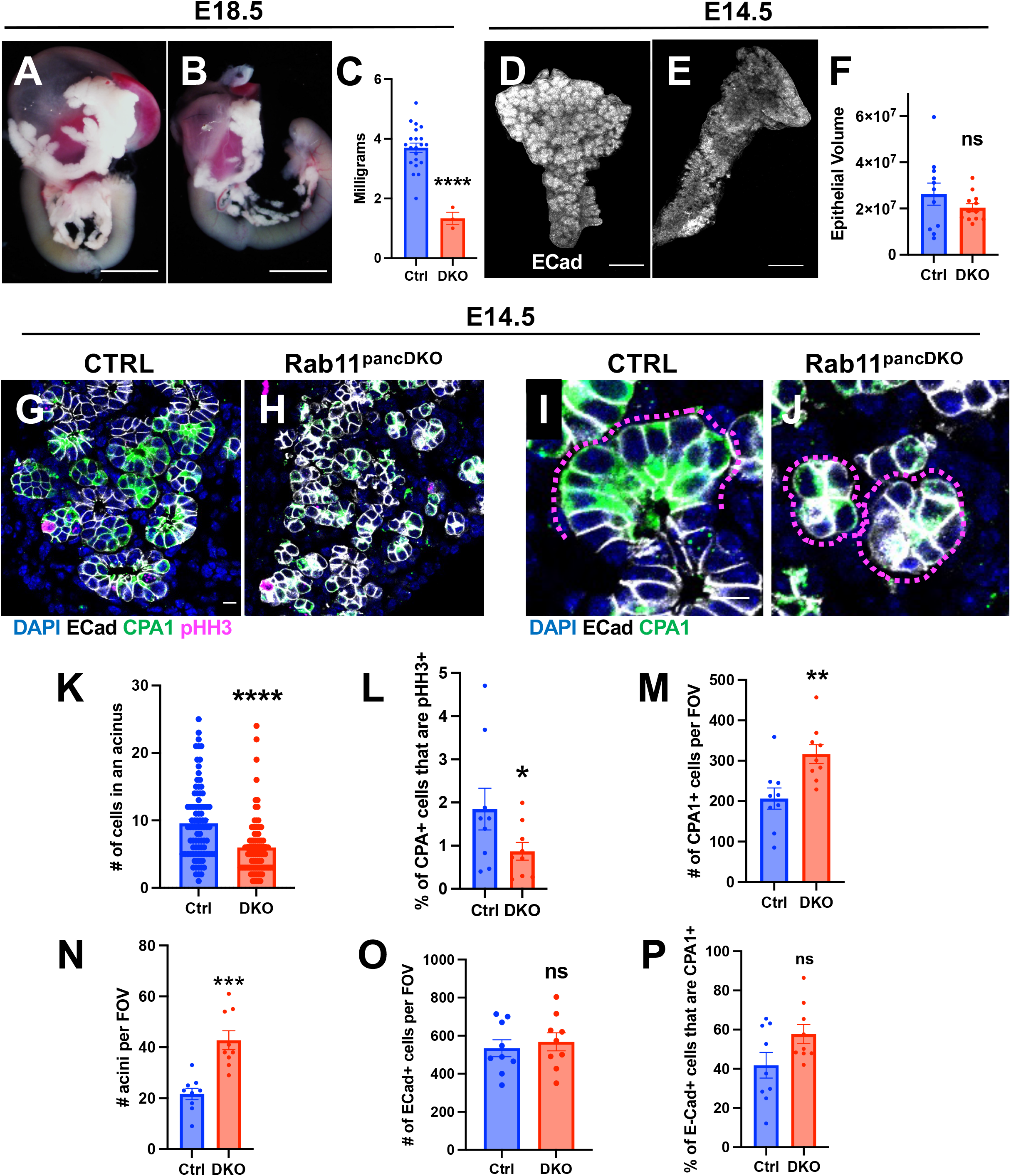
Loss of Rab11 leads to disruption of overall epithelial and acinar morphogenesis. At E18.5, there are obvious morphogenetic defects in the hypoplastic Rab11^pancDKO^ pancreas as compared to a littermate control (A-B), accompanied by a decrease in pancreas weight (C). At E14.5 there is a non-significant trend towards a decrease in epithelial volume (D-F) as shown in whole mount (E-Cad, cyan). In section at E14.5, there are clear morphogenetic defects in the pancreatic epithelium (G-H; E-Cad, white). Upon closer examination, the morphogenesis of acinar cells (CPA1, green) is disrupted (I-J; single acini outlined in magenta). There is a significant decrease in the number of acinar cells per acinus (K) that is associated with a decrease in acinar cell proliferation (G-H, L; pHH3, magenta). Surprisingly, this is associated with an increase in both the number of acinar cells and the number of acini at E14.5 in the Rab11^pancDKO^ pancreata (M-N), while the total number of E-Cad^+^ cells per field of view (FOV) is unchanged (O). Interestingly, the total percentage of E-Cad^+^ cells that are also CPA1^+^ is not significantly increased. n>3 for E18.5, n>=8 for E-Cad volumes, n=3 for CPA1 stains; error bars represent the standard error of the mean; scale bars A-B=0.5cm, D-E=200uM, G-J=20uM,; data analyzed in PRISM by a Student’s t-test.

Given that our primary interest in studying Rab11 was understanding its role in developmental dynamics, we examined developing pancreata at earlier stages. At both E12.5 and E14.5, we observed no externally obvious defects (**Fig. S3 A-D**) and the epithelial volume of the pancreas as measured in IMARIS in Rab11^pancDKO^ and controls were roughly equivalent at E14.5, albeit with a trend towards a decrease in size of the mutant pancreata (**Fig. 3 D-F**). Upon examination of E14.5 mutant pancreata in section, we found that the epithelium and the acini at the tips of branches exhibited clear morphogenetic differences in the mutants (**Fig. 3 G-J**), potentially contributing to the gross branching defects observed at dissection (**Fig. S3 F-I’**). Acinar cells can be recognized by their anatomical location at the tips of ductal branches, which form florets of approximately 8-12 acinar cells around a central lumen, but also by their expression of the acinar cell marker carboxypeptidase A1 (CPA1) (**Fig. 3 G-J**). Mutant acinar cells could also be recognized by the expression of CPA1, however the morphology and organization of these cells were abnormal (**Fig. 3 H, J**).

Unlike acini in controls, the mutants exhibited acini with strikingly fewer cells. Quantification of these structures in the Rab11^pancDKO^ pancreas revealed on average 50% fewer acinar cells (control average =~ 9; Rab11^pancDKO^ average =~ 5) (**Fig. 3 K**), and this was associated with a decrease in acinar cell proliferation in the mutants (**Fig. 3 G-H, L**). Interestingly, both the number of acinar cells and the number of acini per field of view increased despite decreased proliferation rates (**Fig. 3 M, N**). We also found that the density of epithelial tissue was unchanged in the mutants (**Fig. 3 O**). These findings suggest that the Rab11^pancDKO^ epithelium is undergoing aberrant morphogenesis, generating many more, but smaller, acinar tips. Indeed, it is worth pointing out that the overall organ in mutants tends to be slightly smaller, hence the overall number of acinar cells is not likely higher (**Fig. 3 P, Fig. 4 D-F**). Rather, it is likely that the morphogenetic branching of the pancreas is altered in the absence of Rab11.

**Figure 4.**
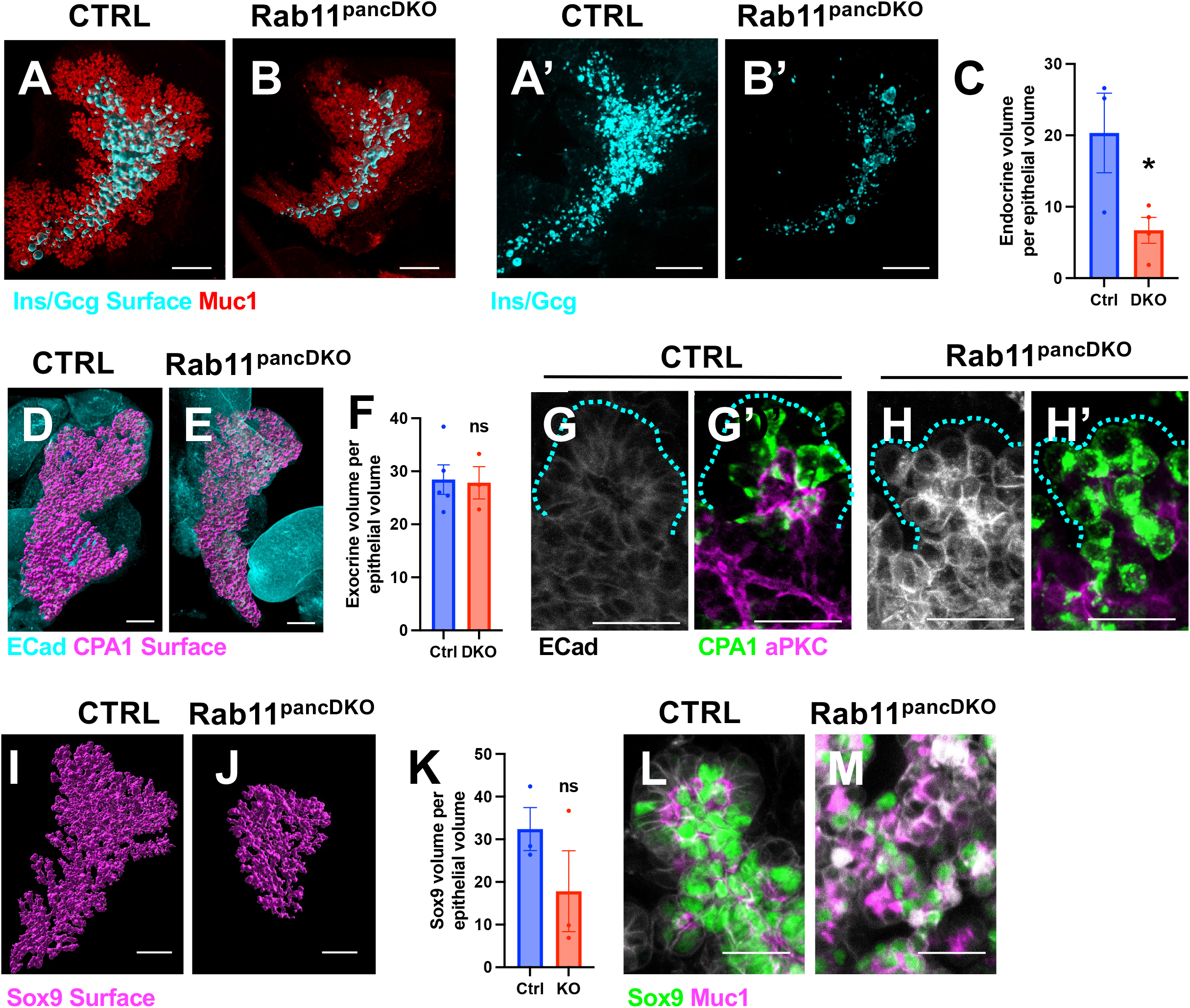
Loss of Rab11 leads to loss of endocrine mass. To assess changes in cell fate determination, E14.5 pancreata were analyzed in whole mount. At E14.5, there is a clear decrease in endocrine volume (insulin/glucagon, cyan; (A, B) IMARIS surface; (A’, B’) whole mount staining). Even when accounting for the trend in epithelial volume decrease in Rab11^pancDKO^ pancreata, there is a statistically significant decrease in endocrine volume (C, uM^3^). In analyzing other pancreatic lineages in a similar way, we found that there is not a detectable change in exocrine volume (CPA1, magenta in (D-F), green in (G’, H’)), although the organization of CPA1 ^+^ cells around lumenal tips (outlined in cyan) is disrupted ((F-G’); E-Cad, white; aPKC, magenta; CPA1, green). We also found that there is a non-significant trend in decrease of ductal Sox9 ((I-J) IMARIS surface, magenta; (L-M) green) in mutant pancreata (I-K). Similar to our findings regarding exocrine cell organization, Sox9^+^ cells that are destined to contribute to a ductal fate are disorganized in the Rab11^pancDKO^ (L-M). n=3, error bars represent the standard error of the mean; scale bars A-B’= 200uM, D-E, I-J=150uM, G-H’, L-M=20uM; data analyzed in PRISM by Student’s t-test.

### Loss of Rab11 leads to loss of endocrine mass

Following our analysis of the morphologically altered acinar lineage in the Rab11^pancDKO^ mutant pancreas, we next asked whether the differentiation of the three main pancreatic lineages was altered. Using whole mount immunofluorescence staining of both control littermate and Rab11^pancDKO^ pancreata, we found that the volume of endocrine cells (Insulin^+^ and Glucagon^+^) per epithelial volume was altogether decreased in the absence of Rab11 (**Fig. 4 A-C**). Consistent with our analysis of CPA1^+^ exocrine cell fate from (**Fig. 3 P**) in section, we did not detect a significant difference between the volume of CPA1^+^ cells per epithelial volume in whole mount (**Fig. 4 D-F**). The organizational defects observed in acini (**Fig. 3 G-J**) were also apparent in whole mount. CPA1^+^ acinar cells are primarily restricted to the end of a lumenal and epithelial branch in controls, but this organization was almost completely disrupted in the mutants (**Fig. 4 G-H’**). In examining ductal cell fate, we did not detect a significant decrease in the volume of Sox9^+^ cells per epithelial volume (**Fig. 4 I-K**), but similar organizational disruption was clearly present in this cell population as well (**Fig. 4 L-M**). Overall, the organization and morphogenesis of the pancreas in the Rab11^pancDKO^ is severely disrupted, and this disruption is correlated with a decrease in endocrine volume. This decrease in endocrine mass may contribute to the defective endocrine function seen in adult mice in (**Fig. 2**).

### Rab11 is necessary for the formation of a lumenal network in the pancreas

Given the known role of Rab11 in epithelial lumen formation (Bryant et al., 2010) and the mislocalization of Rab11 which is observed in lumen-defective Afadin pancreatic mutants (Azizoglu et al., 2017), we sought to understand the possible role of Rab11 in pancreatic lumenogenesis. Using whole mount immunostaining of pancreatic lumens for the lumenal protein Mucin-1 (Muc1), we observed dramatic defects in the formation of the pancreatic lumenal plexus. As early as E14.5, Rab11^pancDKO^ lumens have less distinct boundaries and fail to connect to each other properly (**Fig. 5 A-D**). Indeed, quantifications of the number of disconnected lumens in IMARIS revealed a significant increase in the number of disconnected lumens per total epithelial volume in Rab11^pancDKO^ tissues (**Fig. 5 J**). Furthermore, there are very few obvious ductal structures (**Fig. 5 B, D**). These changes are represented as a model in (**Fig. 5 I**). In control tissues, lumens have well-defined borders and are clearly connected, although it is normal to see some fuzzy lumen-associated protein in some cells. In contrast, the mutant lumens are almost the opposite in that there are very few lumens with well-defined borders, connections are unclear and most of the Muc1 signal has indistinct borders (**Fig. 5 I**).

**Figure 5.**
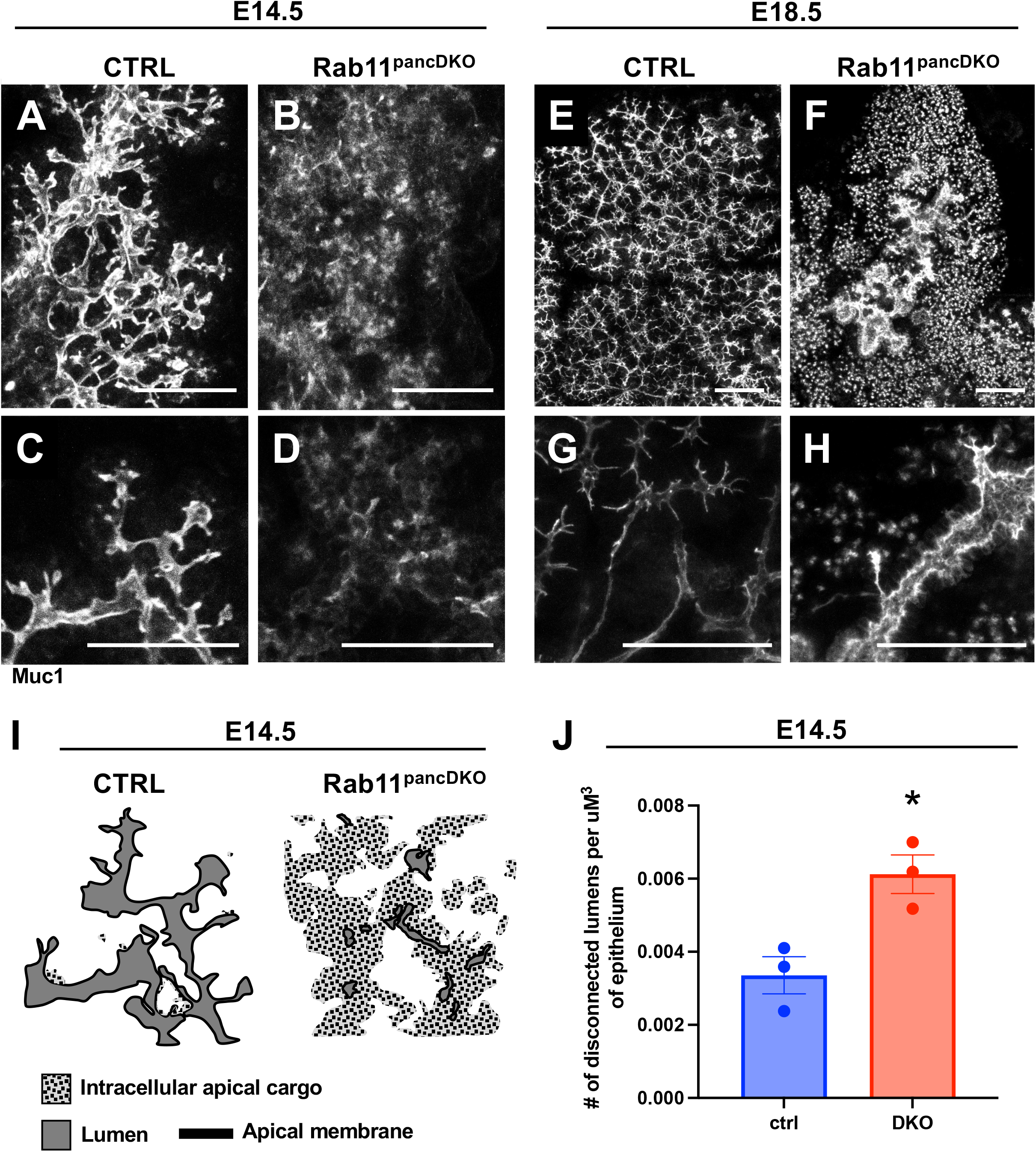
Rab11A & Rab11B are essential for the formation of the pancreatic lumenal plexus. At E14.5, control pancreata have complex lumenal networks as shown in whole mount (Muc1, white) (A). In Rab11^pancDKO^ mutants, maximum intensity projections of the same tissue thickness show indistinct lumen borders and an apparent lack of organization (B). Higher resolution images of thinner stacks show well-connected lumens in control tissues (C), while mutant lumens are largely not connected and display fuzzy, indistinct borders (D). At E18.5, control lumens remain distinct and connected (Muc1, white) (H), while Rab11DKO mutants display a combination of disconnected microlumens and large cyst-like connected lumens (I). Higher resolution images show connected lumens with distinct boundaries in control tissue (J) while the mutant lumens are either small disconnected dots or swollen cyst-like ductal structures (K). The shift away from clear and connected lumens with very few regions of indistinct intracellular cargo is represented in (L), in which lumens are dark grey surrounded by black apical membrane, and intracellular cargo are light grey with black speckles representing apical membrane. Indeed, there is a statistically significant increase in the number of disconnected lumens per total lumen volume at E14.5 (M). n=3 for all experiments; error bars represent the standard error of the mean; scale bars A-D=50uM, E-H=100uM; data in (M) analyzed in PRISM by a Student’s t-test.

At E18.5, we found that the lumens had progressed dramatically in two different directions: small disconnected microlumens or connected but swollen cystic lumens (**Fig. 5 E-H**). Interestingly, the epithelial cells lining these cystic lumens retained expression of Rab11A (**Fig. S4**). This cystic phenotype was particularly surprising given that most of the acini of the pancreas were not connected to these ducts; one might expect smaller, more constricted ducts due to the presumed decrease in fluid volume. Overall, Rab11 clearly plays an integral role in the formation and organization of the lumenal plexus in the developing pancreas.

### Loss of Rab11 leads to defects in secretion of apical cargo

We next assessed Rab11^pancDKO^ pancreata in sections to determine why the mutant lumens were disconnected from each other. Looking more closely at the *Rab11^pancDKO^* mutant tissue at both E14.5 and E18.5, we observed that mutant tissues had severe morphogenetic defects that were associated with disconnected lumens (**Fig. 6 A-D**). Indeed, the failure of lumen continuity appears to derive from a primary failure of lumen formation, both at E14.5 (**Fig. 6 A-B**) and E18.5 (**Fig. 6 C-D**). Upon closer examination of E14.5 Rab11^pancDKO^ mutants, apically-secreted protein Muc1 is seen trapped inside Rab11^pancDKO^ mutant cells, resulting in the indistinct lumen boundaries observed in whole mount (**Fig. 4 E-E’, F-F’**). We also noted that adherens junction protein E-Cadherin (E-Cad) localization shifts upon loss of Rab11, resulting in an increase in intracellular retention of E-Cad (**Fig. 6 E’’, F’’**). In addition, we note its aberrant retention at the AMIS marked by Muc1. Furthermore Podocalyxin (Podxl), a protein that both marks and facilitates the formation of lumens, is similarly either trapped inside cells or completely undetectable (**Fig. 6 G-H’’**). These findings suggest that Rab11 is critically required for faithful trafficking of required components during AMIS formation.

**Figure 6.**
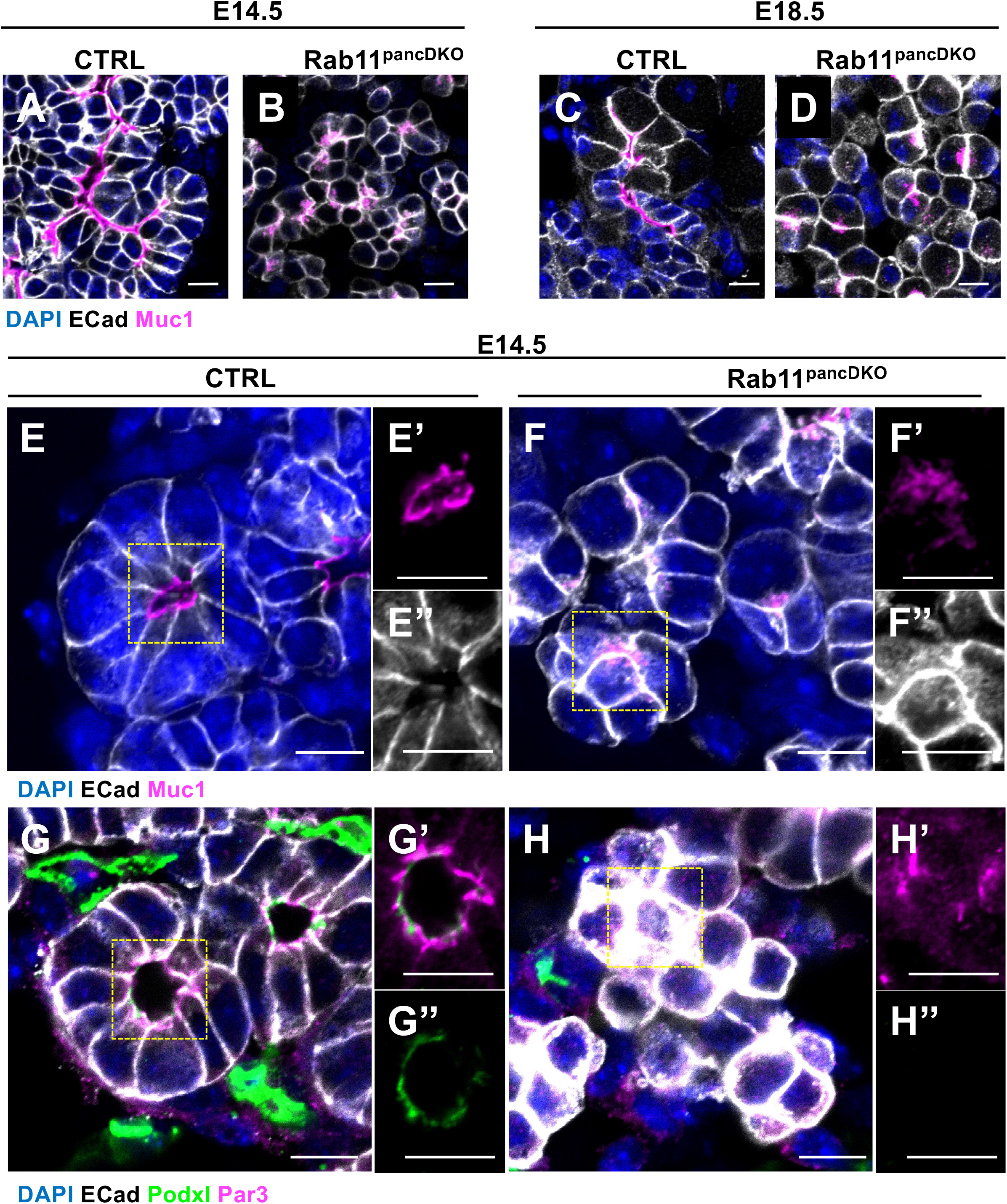
Loss of Rab11 leads to retention of apically-targeted cargo. Sections through tissue at both E14.5 (A, B) and E18.5 (C, D) show changes in epithelial morphology (E-Cad, white) as well as lumen connectivity (Muc1, magenta; DAPI, blue). Upon closer examination, E14.5 control tissue displays clear open lumens (Muc1, magenta) with distinct boundaries (E) and an apical membrane region (E’) that largely excludes E-Cad (white) (E’’). Rab11^pancDKO^ tissue instead shows intracellular retention of lumen marker Muc1 (magenta) that correlates with a failure of lumen formation (F, F’). Trafficking failures are further confirmed by the strong presence of E-Cad around each cell border, even where lumen marker Muc1 is localized (F’’). Additionally, there is an increase in intracellular E-Cad (F’’, yellow arrowhead). Lumen marker Podxl (green) is similarly localized to the apical membrane (Par3, magenta) in open lumens in control tissue (G-G’’), but is strongly reduced or completely missing from Rab11^pancDKO^ tissue (H-H’’). n=4 for A-D; n>3 for E-F’’; n=3 for G-H’’; scale bars=10uM, yellow boxes indicate the location of insets for the following panels.

### Rab11^pancDKO^ cells exhibit abnormal cell polarization

Given that proper transport of cellular elements is critical to establishment and maintenance of cell polarity (Roman-Fernandez and Bryant, 2016), we further investigated the localization apical and basal components in *Rab11^pancDKO^* epithelial cells. As shown in (**Fig. 6**), we had noted intracellular retention of Muc1. We also found that there was mislocalization of Muc1 to the lateral or even the basal membrane in some cells (**Fig. 7 A-B’**, white asterisks). Furthermore, some mutant cells seemed to make more than one AMIS, thus participating in more than one attempt to form lumens between neighbors (**Fig. 7 B’**, yellow arrowhead).

**Figure 7.**
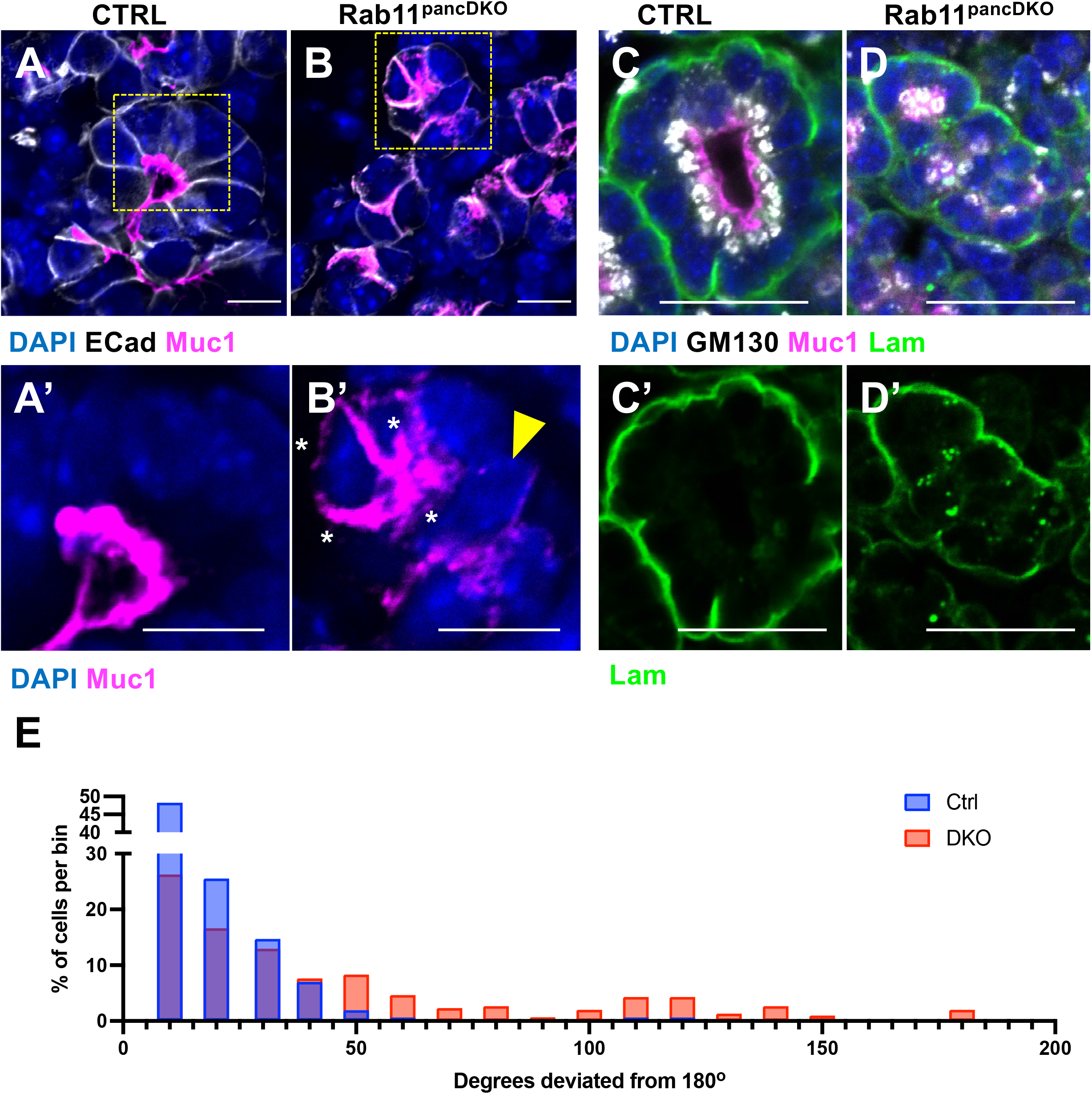
Rab11^pancDKO^ cells exhibit abnormal cell polarization. In contrast to control tissue where most cells have a single apical membrane (A, yellow box; A’), Rab11^pancDKO^ mutant cells were observed to have apical markers (Muc1, magenta) on multiple membranes (B, yellow box; B’, white asterisks) and/or participate in multiple apical sites (B’, yellow arrowhead). To assess overall cell polarity, control and Rab11^pancDKO^ tissues were stained for basement membrane (Laminin, green), nuclei (DAPI, blue), Golgi (GM130, white) and apical membrane (Muc1, magenta) (C-D). The angles created by lines drawn from basement membrane to Golgi (through the nucleus) and from the Golgi to the apical membrane were measured in FIJI. The deviation of each measurement from the “perfectly” polarized cell represented by a 180° angle was calculated and binned into 10° deviation categories. Plotting the % of total cells per bin revealed a clear shift in cell polarity (E). Of note, there also seems to be defects in trafficking Laminin in mutant cells (D’). n>3 for A-B’; n=3 for C-E; scale bars=10uM.

This observation led us to examine the overall polarity of these cells by staining E14.5 sections for Laminin (basal extracellular matrix), GM130 (Golgi), Muc1 (apical membrane) and DAPI (nucleus) (**Fig. 7 C-D’**). We found that Rab11^pancDKO^ cells displayed both punctate internal laminin staining (**Fig. 7 D’**), as well as abnormally localized GM130 (**Fig. 7 C-D, E**). To quantify potential defects in cell polarity, a line was drawn in FIJI from the basal membrane through the nucleus to the Golgi, and another line was drawn from the Golgi to the apical membrane. The angle formed by these two lines was quantified and compared between controls and mutants, with the assumption that an angle of 180° (a straight line) would represent a perfectly polarized cell. There was a clear shift in the distribution of polarity angles away from 180 ° in the mutant cells, with some cells even displaying angles that are never seen in control cells (including 0°, indicating cells whose basal and apical membranes overlap) (**Fig. 7 E**). Overall, Rab11^pancDKO^ cells display severely altered cell polarity.

### Rab11 is required for the coordination of a single AMIS

We next asked whether polarity-determining proteins such as members of the Par (Par6b, Par3, aPKC) or Crumbs (Crbs3) complexes were mislocalized in the Rab11^pancDKO^ cells, as has been reported previously. Indeed, although a portion of each polarity protein assayed was able to incorporate into an AMIS, there was an increase in intracellular signal from each protein, as well as a significant increase in the number of apical sites per cell in the mutants (**Fig. 8 A-H’, K, L**). This reflects the localization of apically-targeted proteins to multiple regions of the cell membrane shown in (**Fig. 7 B-B’**). Tight junction protein ZO-1 largely followed the localization of apical membrane in Rab11^pancDKO^ cells, but we also observed instances where there were tight junctions both within and between cells where there was no apical membrane present (**Fig. 8 I-L**). These data suggest that Rab11 is critical for the coordination of a single AMIS between groups of cells by affecting the localization of polarity-determining proteins. Our findings from (**Fig. 7**) and (**Fig. 8**) are summarized in (**Fig. 8 K**), where in the mutant cells the Golgi (grey) is improperly oriented, multiple AMIS’ (green) are forming, junctions (red) are mistargeted, and both apical (green) and basal (magenta) components are intracellularly retained. Given these observations, it is no surprise that pancreatic morphogenesis is so severely disrupted in the Rab11^pancDKO^. Altogether, our results suggest that Rab11 regulates initial molecular composition of the lumen initiation site (AMIS), and is therefore essential for overall pancreas morphogenesis and function via its crucial role in lumen formation.

**Figure 8.**
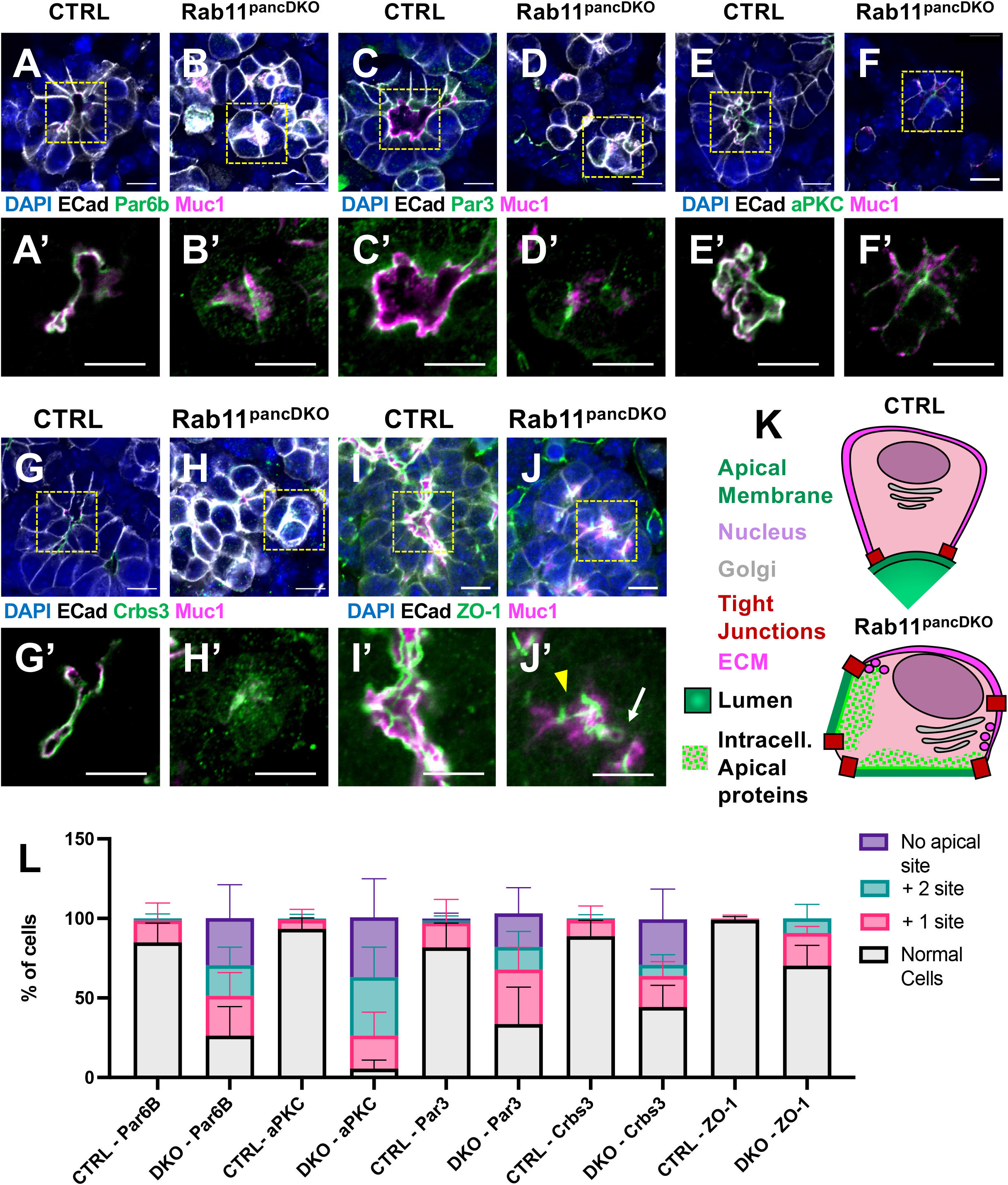
Rab11 is required for the coordination of a single apical membrane initiation site. Localization of apical polarity determining Par complex (A-F’) and Crumbs complex (G-H’) markers were assessed in control and Rab11^pancDKO^ tissue. Par6b (A-B’), Par3 (C-D’), and aPKC (E-F’) (green) localized tightly to the apical membrane (Muc1, magenta) in control tissue. In Rab11^pancDKO^ tissue, a fraction of each of these Par complex components were able to properly localize to the apical membrane, but many mutant cells displayed intracellularly-retained polarity markers as well as the formation of multiple apical sites per cell (B’, D’, F’). Similarly, Crumbs complex member Crumbs3 (G-H, green) is tightly localized to the apical membrane (Muc1, magenta) in control tissue (G-G’), while intracellular retention of Crbs3 is increased in the Rab11^pancDKO^ (H-H). Furthermore, tight junction marker ZO-1 (magenta) localizes to cell-cell contacts beneath the apical membrane (aPKC, green) in control tissue (I-I’). In mutant tissue, ZO-1 continues to associate with regions of apical membranes, but is also ectopically present within (J’, white arrow) or between (J’, yellow arrowhead) cells that are not coordinating an AMIS (J-J’). A model summarizing the defects found in Fig. 7 and Fig. 8 is shown in (K) (cell, pink; laminin, magenta; nucleus, periwinkle; Golgi, lavender; ZO1, cyan; apical polarity determining complex proteins, green; lumen, grey). The percent of cells that have ectopic apical sites as shown by Par6b, Par3, aPKC, Crbs3 or ZO-1 staining are quantified in (L). There is no significant difference (p>0.05) between the percentage of cells with 1 extra Par6B site, with 1 extra aPKC site, with 2 extra Par3 sites, or with 1 or 2 extra Crbs3 sites. There is also no significant difference between any ZO-1 measurements. There is an extremely significant difference (0.001>p>0.0001) between the percentage of cells with 2 extra Par6B sites and with no Par3 sites. There is an extremely significant difference (p<0.0001) between the percentage of cells with a single Par6B site, with no Par6B site, with a single aPKC site, with 2 extra aPKC sites, with no aPKC site, with a single Par3 site, with a single Crbs3 site, and with no Crbs3 site. N=3; scale bars=10uM; data in (L) were analyzed by a 2-way ANOVA with multiple comparisons in PRISM.

## DISCUSSION

Here we report that loss of Rab11A and Rab11B (Rab11^pancDKO^) in the developing mouse pancreas results in defects in pancreatic morphogenesis that are linked to a failure of singular AMIS and single lumen formation. This is a novel finding because AMIS formation and subsequent single lumen formation has primarily been studied in *in vitro* systems in which lumens are being formed between two cells. In the developing pancreas and in other developing organs, lumens are instead formed between groups of four or more cells. Previous models of *de novo* lumen formation have not taken into account the unique organizational needs of larger groups of cells. In the absence of Rab11, cells form multiple AMISs and fail to coordinately form a single central lumen. As a result, the pancreatic epithelium fails to establish the connected lumenal plexus. These lumen formation defects are linked to dramatic changes in pancreatic morphogenesis. The Rab11^pancDKO^ pancreata also display a decrease in endocrine cell mass, which is likely linked to the defects in ductal plexus formation and morphogenesis (Azizoglu et al., 2017). This study underscores the stark requirement for Rab11 during pancreas formation, and specifically show that Rab11 is essential for epithelial lumen formation during organogenesis.

Our work, together with decades of in *vitro* molecular studies of Rab11 and vesicle trafficking in general (Ren et al., 1998; Stenmark and Olkkonen, 2001) allow us to speculate as to why loss of Rab11 may so profoundly impair murine pancreas development. The primary molecular function of Rab11 is to facilitate recycling of various cargoes around the cell. It has been shown by others that Rab11 helps vesicles bud off the recycling endosome and targets them to the correct region of the plasma membrane via coordination with Rab8 and Myosin5B (Bryant et al., 2010; Hales et al., 2002; Roland et al., 2011). This process has been closely studied at the apical membrane, but Rab11 is also known to traffic cargoes such as E-Cadherin and integrins to and from the lateral and basal membranes, respectively (Desclozeaux et al., 2008; Moreno et al., 2022; Tanasic et al., 2022; Woichansky et al., 2016). Indeed, we speculate that Rab11 recycling of lateral membrane cargo to the apical membrane, as well as its role in promoting apical membrane turnover, are both critical to lumen formation (**Fig. 9**).

**Figure 9.**
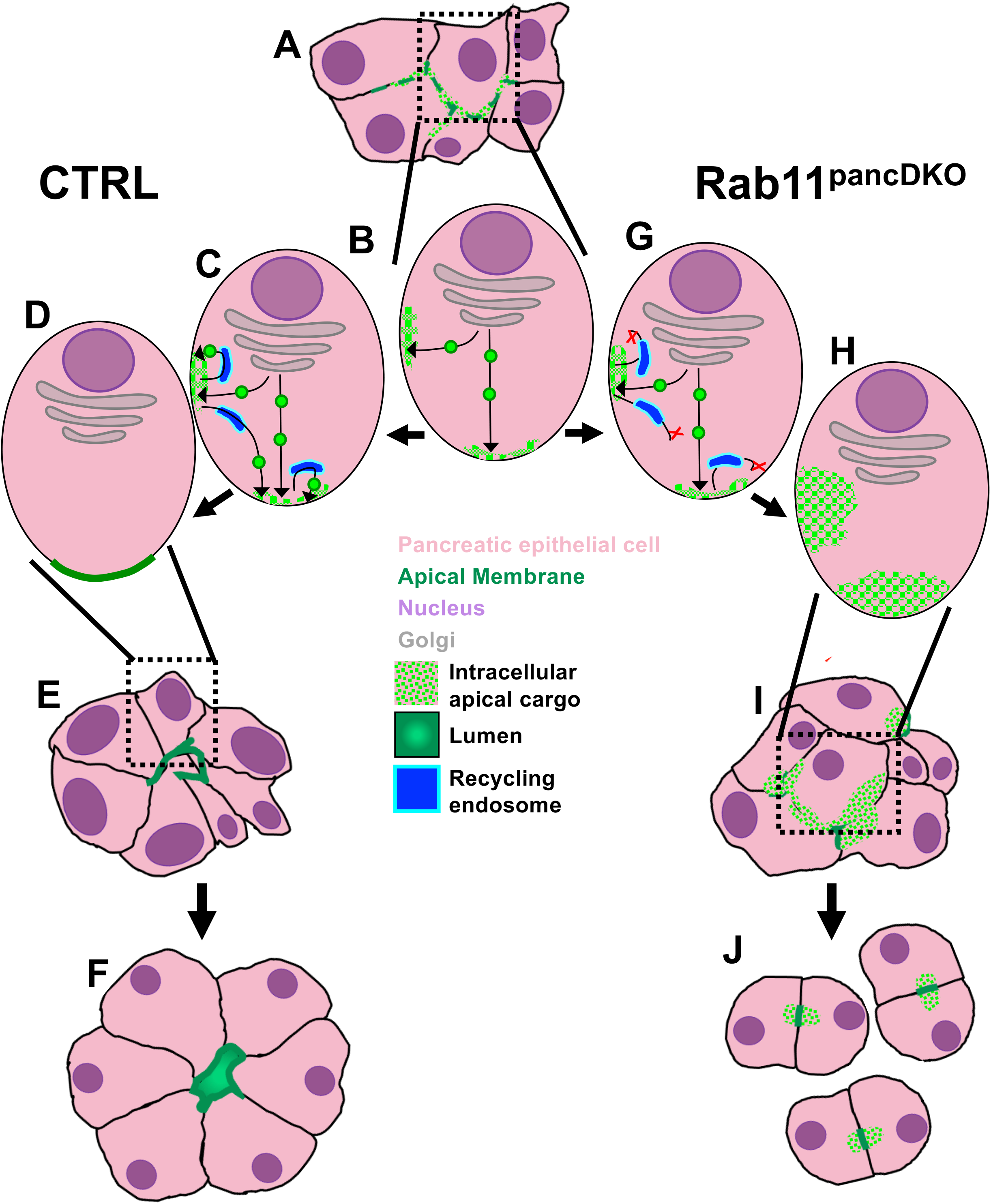
Model: Rab11 directs the formation of a singular apical membrane initiation site through the proper membrane-localization of apical and junctional proteins. During early stages of pancreatic development, cells have not yet coordinated a single AMIS and there is often more intracellular apical cargo even in wild-type cells compared to later stages of pancreas development (A). Based on the known roles and functions of Rab11 from other studies, we propose that in the pancreatic epithelium Rab11 directs and enables recycling of apical and junctional components (B). In particular, we propose that Rab11 regulates normal turnover of membrane-associated proteins at both the lateral and apical membranes (this could also be true of basal proteins, but our study focused on the apical side of the cell), as well as the re-localization of proteins from the lateral membrane to the apical membrane (B), thus orchestrating a single AMIS (C). Pancreatic epithelial cells continue to traffic and exocytose various proteins, enabling the cells to open a single lumen between themselves (D). We propose that the redirection of recycling membrane is disrupted in our Rab11^pancDKO^ cells (E). Apical and junctional proteins are still trafficked to the apical and lateral membranes, but after uptake of membrane for either normal turnover or lateral-to-apical re-localization, proteins are unable to return to the membrane in the absence of Rab11 (E). This leads to the formation of multiple apical sites per cell and a failure to clear adherens junction protein E-Cad from the apical membrane (F). Mysteriously, midgestational pancreatic cells are able to somehow shift from multicellular blobs to primarily twocell structures surrounding a single AMIS whose necessary apical proteins are trapped inside each cell (G). Pancreatic epithelial cells, pink; apical membrane, green; nucleus, purple; Golgi, grey; intracellular apical cargo, bright green spots on a pink background; lumen, green; recycling endosomes, blue.

Together, our findings in this study led us to propose the following model: Rab11 normally drives regular turnover at both the lateral and apical membranes, as well as redirection of proteins from the lateral to the apical membrane. Tight regulation of these trafficking movements allows cells to coordinate in the formation of a single AMIS (**Fig. 9 A-D**) that then expands into an open lumen (**Fig. 9 D-F**). In the Rab11^pancDKO^ pancreatic epithelium, cells are capable of initiating polarity formation, AMIS formation and lumen formation (**Fig. 9 B, G**), however we speculate that loss of Rab11 blocks the redirection of membrane and proteins out of the recycling endosome back to the plasma membrane, resulting in cells that are unable to drive lumenogenesis across the finish line (**Fig. 9 G, H**). We propose that new proteins and membrane are continually synthesized and trafficked to the plasma membrane and constantly turned over and sorted through the recycling endosomes, but they are unable to be properly targeted back to the plasma membrane in the absence of Rab11 (**Fig. 9 G, H**). This phenomenon may drive the intracellular buildup of apical and junctional cargo in the Rab11^pancDKO^, rather than a direct failure of initial targeting and exocytosis.

Our findings that multiple apical sites form in the absence of recycling regulator Rab11 suggest that cells may normally initially target apical components to more than one location in the cell during AMIS formation, only to be “cleaned up” and recycled away from the forming lateral membrane to the apical membrane (**Fig. 9 C, D**). While this hypothesis requires further testing, it may shed some light on how cells accomplish the feat of choosing and coordinating an AMIS in large groups of cells *in vivo* (**Fig. 9 A-F**). For indeed, in the Rab11^pancDKO^ (**Fig. 9 F**) and in Rab11 knockdown or mutation studies in other systems (Bryant et al., 2010), groups of cells form multiple lumens (or attempt to, in the case of the Rab11^pancDKO^). How cells in the Rab11^pancDKO^ are able to largely overcome this multiple-lumen initiation over the course of development, and remodel into pairs of cells with stunted AMIS-like foci and no lumen (**Fig. 9 I-J**), requires further study. Future studies of this mouse model will help us better understand pancreatic morphogenesis and how it interfaces with lumen formation.

Given our observations, we propose pancreatic microlumens form similarly to Madin-Darby Canine Kidney (MDCK) cell lumens, a process extensively studied by the Mostov group and its scientific descendants (Bryant et al., 2010; Flasse et al., 2021; Kim et al., 2015; Mostov and Deitcher, 1986; Pollack et al., 1997; Pollack et al., 1998; Roland et al., 2011). Findings from our lab (Villasenor et al., 2010) and others (Roman-Fernandez and Bryant, 2016) show that, similar to MDCK cells, junctions and apical polarity determining complex proteins, such as the Par complex and the Crumbs complex, are coordinately targeted to a single apical location that is “agreed upon” by all the pancreatic epithelial cells participating in the newly forming lumen. These cells form a three dimensional (3D) rosette of polarized cells that initiate single lumen formation at their center. The opening of a microlumen is facilitated in part by secretion of negatively-charged Podocalyxin, creating ionic repulsion forces, as well as by actomyosin-generated contractile forces (Camelo and Luschnig, 2021; Dekan et al., 1991; Schottenfeld-Roames et al., 2014; Strilic et al., 2009). Once a pancreatic microlumen has been formed through these mechanisms, it then fuses with other microlumens and with the foregut lumen (Villasenor et al., 2010). Our work shows that these events are mediated by Rab11.

Another open question in the field focuses on how junctions are cleared from the AMIS to facilitate detachment of cells during lumen formation. Two possible mechanisms have been proposed: passive or active junction clearance. In the passive junction clearance model, junctional proteins are pushed to the periphery by continuous addition of new membrane lacking junctional proteins to the AMIS. While it is known that there is a switch in apical membrane biogenesis in which junction proteins are no longer added, simply displacing junction proteins from the AMIS may not fully account for their continued restriction to the periphery (Marciano, 2017). In the active model, junction proteins are endocytosed from the AMIS and recycled to the periphery (Flasse et al., 2021; Yan et al., 2016). Our findings that E-Cadherin is retained at the apical membrane upon loss of Rab11 suggests that recycling is indeed required to clear junctions from the apical membrane to facilitate lumen opening. This model aligns with active clearance of junctions observed in our previous studies in blood vessel lumen formation that identify active clearance of junctions (Barry et al., 2016). Altogether, we favor an active role for apical membrane clearance in which Rab11 plays an essential role.

Our data underscore the direct link between epithelial morphogenesis, architecture and cell fate. In our previous studies, we showed that when the pancreatic epithelium fails to transform from a stratified epithelium to a ramifying tree of continuous ducts containing functional lumens, the emergence of endocrine cells is altered (Azizoglu et al., 2017). In that study, the loss of the protein Afadin and its partner RhoA led to a delay in the resolution of the lumenal plexus, which we postulated resulted in additional cycles of proliferation of endocrine progenitors. Given that Rab11 was mislocalized in the absence of Afadin, we predicted the ablation of Rab11 would yield a similar phenotype. However, in the absence of Rab11 we observe instead the loss of endocrine cells. We ascribe this difference to the different anatomy of the mutant epithelium, whereby rather than large, multi-layered epithelial aggregates observed in Afadin^pancDKO^, we find a breakup of the epithelium into finer structures in the Rab11^pancDKO^ (as shown by cross sections showing 2-cell wide strands, **Fig. 6 A-D**). These observations support the existence of a 3D niche for endocrine development that depends on proper lumen formation.

Additionally, it is worth noting that the exocrine machinery of the pancreas similarly depends on faithful epithelial morphogenesis and lumen formation. Our data show that normal acinar and ductal fate is tied to coordination of epithelial cells to form pre-acinar tips of a certain size. Loss of Rab11 leads to smaller tips and disconnected ducts. These defects continue to be present after birth, where viability of pups is impaired. Whether postnatal survival is primarily driven by the decrease in endocrine mass observed at E14.5, or by the likely decrease in digestive enzyme secretion resulting from a disconnected pancreatic tree is unclear, but we speculate that both are at least partially responsible.

Altogether, our study provides novel insights into the mysteries of pancreatic morphogenesis and lumen formation. We posit that loss of Rab11 blocks vesicles from budding off recycling endosomes, resulting in failure of directional trafficking within epithelial cells, and defects in cell polarity (demonstrated by multiple apical membrane initiation sites per cell) and junction remodeling. These cellular changes, in turn, result in failure of lumen formation and connection, as well as severely disrupted epithelial morphogenesis. This study opens the door to closer scrutiny of pancreatic morphogenesis, especially given the building link between developmental morphogenesis and disease. In the future, we hope to utilize this mutant and others to build a complete picture of pancreatic morphogenesis and to continue teasing apart the links between lumen formation and morphogenesis.

## MATERIALS AND METHODS

### Mice & Embryo Handling

All animal husbandry was performed in accordance with protocols approved by the University of Texas Southwestern Medical Center Institutional Animal Care and Use Committee. E10.5-E18.5 embryos and postnatal tissues were collected and dissected in PBS. Tissues and embryos were photographed at dissection with a ZEISS NeoLumar S 0.8x FWD 80mm Microscope, and embryos, pups and adults were photographed with an iPhone12. Tissues were fixed in 4% paraformaldehyde (PFA) in PBS overnight at 4°C. CD1 or mixed-background littermates were used as controls. Rab11A^f/f^, Rab11B^-/-^, and Pdx1-Cre lines were used in this study (see references cited herein). We originally attempted to generate Rab11^pancDKO^ mutants by crossing Rab11A^f/f^;Rab11B^-/-^ females with Rab11A^f/f^;Rab11B^-/-^;Pdx1-Cre males. While the few surviving Rab11^pancDKO^ males were viable and fertile, the defects associated with these mutations applied selective pressure and blocked the effective recombination of the Rab11A flox sites by the Cre. Subsequently, males were maintained as Rab11A^f/+^;Rab11B^-/-^;Pdx1-Cre or Rab11A^f/f^;Rab11B^+/-^; Pdx1-Cre and bred to Rab11A^f/f^;Rab11B-/- females to generate Rab11^pancDKO^ tissue.

To monitor survival in postnatal mutants, cages with plugged females were monitored from E18.5 to what would be E22.5 for the birth of a new litter. Due to a high frequency of cannibalism, the number of pups was estimated at P0 with minimal perturbations to the cage. Pups were then counted and photographed every day until P8 to track any deaths or failures to thrive. If dead pups were observed in the cage, tissue was collected for genotyping. Toes from live pups were collected for genotyping between approximately P5 and P9, and then mice were weaned between P21 and P28. At weaning, mice were weighed in a cup using a Kitchen Tour Digital Touch scale and photographed using an iPhone 12.

### Glucose tolerance assay

To test the glucose tolerance of adult mice, mice were fasted for at least 12 hours (overnight). Mice were then weighed and their glucose (D+ glucose, Sigma G8270) dose was calculated (2mg glucose / g body weight). A baseline blood glucose reading was taken using a True Balance glucometer with True Balance test strips. A fresh razor blade was used to cut off a small piece of the end of the tail, and blood was collected as a bolus that was directly applied to the test strip. Mice were then injected with glucose intraperitoneally, and their blood glucose levels were measured at 15, 30, 60 and 120 minutes. These values were plotted in PRISM and analyzed by 2way ANOVA.

### Immunofluorescence on sections

After fixation, tissues were rinsed thrice in PBS then dehydrated via an ethanol gradient and washed twice for 30min in 100% ethanol. Either immediately or after storage at −20°C, tissues were cleared by washing in xylene twice for 10min and then transitioned into paraplast (McCormick Scientific) by washing in 1:1 xylene:paraplast for 10min at 60°C. Tissues were subsequently washed approximately once per hour for at least three hours and soaked overnight in paraplast at 37°C. Tissues were embedded in paraplast and sectioned at either 10uM or 20uM with a Biocut 2030 microtome or a HistoCore MULTICUT semi-automated rotary microtome onto SuperfrostPlus glass slides (Fisher).

As previously described (Azizoglu et al, 2017), sectioned tissue was deparaffinized with xylene and then rehydrated through an ethanol series. Sections were permeabilized for 10min in 0.3% Triton X-100 in PBS then subjected to antigen retrieval with either R buffer A (nuclear antigens) or R buffer B (cytoplasmic antigens) in a 2100 Retriever (Electron Microscopy Services). Slides were blocked with 5% Normal Donkey Serum (NDS) in PBS for two hours, then were incubated in primary antibody diluted in 5% NDS overnight at 4°C (**Table S1**). The next day, slides were washed in PBS and incubated in secondary antibody diluted in 5% NDS for two hours (all secondary antibodies were AlexaFluor). Slides were subsequently washed in PBS and blood cells were lysed by a 15min incubation in 10mM CuSO_4_, 50mM NH_4_Cl @ pH=5 solution, followed by a 5min wash in H_2_O. Finally, slides were washed in PBS and mounted with DAPI Fluoromount-G (Southern Biotech). Stained slides were imaged on a Nikon A1R confocal microscope provided by the UT Southwestern Molecular Biology Department.

For antigens requiring additional antigen retrieval, Tyramide signal amplification was performed in accordance with the manufacturer’s protocol (Life Technologies) and as previously described (Braitsch et al., 2019).

### Whole-mount Immunofluorescence

As previously described (Daniel et al., 2018), fixed tissues were dehydrated to 100% methanol via a gradient and stored at −20degreesC until use or transferred directly from PBS to 100% ice cold methanol for a secondary fixation for 30min at room temperature (RT) and then washed back to PBS via a gradient. Once rehydrated, tissues were permeabilized for 1.5-2 hours for tissue between E10.5 and E12.5, 3 hours for E14.5 and 4 hours for E18.5 tissues and then blocked in CAS Block (ThermoFisher) for at least two hours. Samples were incubated in primary antibody diluted 1:500 in CAS Block overnight at 4°C and were washed in PBS at RT for at least six hours the next day before incubating in secondary antibody diluted 1:500 in CAS Block overnight at 4°C. Tissues were then dehydrated into 100% methanol and cleared in a 1:2 mixture of benzyl alcohol/benzyl benzoate (BABB) for at least ten minutes at RT. Samples in BABB were visualized in concave glass slides using an LSM710 Meta Zeiss confocal provided by the UT Southwestern Molecular Biology Department.

### Cell polarity quantifications

Cell polarity was quantified on E14.5 tissue sections stained with Laminin (basement membrane), DAPI (nuclei), GM130 (Golgi marker) and an apical marker (often Muc1). Cells were only measured if they were clearly a part of an acinar tip (ductal cells were excluded to the best of our abilities). The “Angle” tool in Fiji was used to manually draw an angle from the laminin to the GM130 signal (passing through the nucleus), and then from GM130 to the apical marker. The starting point on the basement membrane was selected as the region aligned with the center of the nucleus, and the line drawn from the laminin to the GM130 created an approximate 90° angle from the line to the laminin. Similarly, the line from GM130 to the apical membrane was drawn so the line contacted the closest point of the apical membrane at an approximate 90° angle. If a single cell had more than one apical membrane site, the measurements were repeated for each apical membrane site starting from the same laminin and GM130 locations. The raw values were then subtracted from 180degrees (the perfect ideal for a polarized cell) and the absolute value was calculated. Values were manually binned into 10° bins for ease of data representation in Excel. The degree of variation from 180° was then plotted and analyzed in via 2-way ANOVA in PRISM.

### Cell-cell junction and polarity marker quantifications

The number of concentrated patches of junction markers (ZO-1) and polarity markers (Par3, Par6b, aPKC, Crumbs3) were manually counted in individual pancreatic epithelial cells (E-Cad). The expectation for a perfectly polarized cell was used as the baseline for comparison – for junction markers, there should be two punctae of strong signal on either side of an apical membrane patch, while for apical markers there should only be one region of strong signal per cell on the plasma membrane. These counts were then displayed as the number of ectopic junctional or apical sites per single control or DKO cell. Results were graphed and analyzed via ANOVA in Prism.

### Lumen discontinuity quantifications

Whole-mount immunofluorescent images of pancreata stained for E-Cad and Muc1 were loaded into the image analysis program IMARIS. A surface of E-Cad (detail level 10uM to reflect the approximate size of a single cell) was created to quantify the total volume of each pancreas. A surface of Muc1 (detail level 2.5uM) was created, and then the largest volume object (the primary central plexus of connected lumens) was deleted. The number and volume of lumens disconnected from the central plexus were quantified. These values were normalized to the volume of each pancreas in Excel and then plotted and analyzed by Student’s T-Test in PRISM.

### Pancreatic lineage volume quantifications

Whole-mount immunofluorescent images of pancreata stained for E-Cad and a lineage marker (Insulin & Glucagon; CPA1; Sox9) were loaded into the image analysis program IMARIS. A surface of a manually-determined Region of Interest (ROI) of E-Cad signal (detail level 2.5uM) was created to quantify the total volume of each pancreas (other parameters were determined automatically by IMARIS). Dorsal pancreas only was measured, with the cutoff point for ROI determination being where the tissue is most narrow before expanding into the ventral pancreas. Another surface of the lineage marker (detail level 2.5uM) using the same ROI was created to quantify the volume of that cell population within the pancreas (other parameters were determined automatically by IMARIS).The sum of the volume measurements for each surface were recorded in excel. Lineage volumes were standardized to the total volume of the individual epithelium, and those values were plotted and analyzed by Student’s t-test in PRISM.

### Number of cells per field of view quantification

To quantify morphological changes in the acinar cell compartment, cells as viewed by immunofluorescence in section were determined to be of acinar fate by CPA1 staining. Using the “Cell Counter” function in FIJI, the number of cells per acinar cluster at E14.5 and the total number of CPA1^+^/pHH3^+^ cells, CPA1^+^ cells, pHH3^+^ cells, E-Cad^+^ cells, and acini were quantified. This was done for all fully visible acinar rosettes in at least three fields of view of three sections of each biological replicate. These values were plotted and analyzed by Student’s T-Test in PRISM.

### Data analysis and visualization

Data were both analyzed and plotted in PRISM 9 XML. Statistical tests (indicated in the relevant methods sections and figure legends) utilized were the parametric unpaired Student’s t-test and the 2-Way ANOVA with Tukey’s multiple comparisons test (“compare cell means regardless of rows and columns”). ns=p>0.05; *=0.01<p<0.05; **=0.001<p<0.01; ***=0.0001<p<0.001; ****=p<0.0001. Images were manipulated in FIJI. Figures and models were made using Microsoft Powerpoint, data were partially analyzed in Microsoft Excel and text was written in Microsoft Word.

## ACKNOWLEGEMENTS

We would like to thank the entire Cleaver lab, as well as Drs. Elizabeth Chen, Michael Dellinger, Caitlin Maynard and Neal Alto, for discussion and critiques throughout this study.

## Author contributions

Conceptualization: H.R.B., D.B.A., D.M., O.C.; Methodology: H.R.B., D.M., O.C.; Validation: H.R.B., T.B., L.F.; Investigation: H.R.B.; Resources: H.R.B., N.G., O.C.; Data curation: H.R.B., D.M., O.C.; Writing – original draft: H.R.B., O.C.; Writing – review and editing: H.R.B., D.M., N.A., T.B., N.G, O.C.; Visualization: H.R.B.; Supervision: O.C., D.M.; Project administration: O.C.; Funding acquisition: O.C., H.R.B.

## Supplementary information

Supplementary information available online at …

