## Supplemental figure legends and tables for "Rab11 is essential to pancreas morphogenesis, lumen formation and endocrine mass"

### SUPPLEMENTARY MATERIALS

### SUPPLEMENTARY FIGURE LEGENDS

**Figure S1 – Rab11 is expressed in all pancreatic lineages at E18.5.** As shown by immunostaining of sectioned control tissues, Rab11A (green) is enriched at the apical membrane of CPA1$^+$ (white) acinar cells (A-A'), as well as at the apical membrane of Sox9$^+$ (white) ductal cells (B-B') at E18.5. Differential staining techniques utilized to reveal nuclear antigens yielded lateral enrichment of Rab11A in in B-B', but this enrichment was not observed using our standard staining methods. Rab11A is also lowly expressed in the Insulin$^+$ and Glucagon$^+$ (white) endocrine population (C-C'), as shown by the yellow arrowheads. A representative section through a control E18.5 pancreas (D-D') shows that Rab11A (green) is expressed throughout the pancreatic epithelium (E-Cad, white) but is absent from some surrounding mesenchymal cell populations. Regions outside the pancreatic epithelium that show high expression of Rab11A are likely blood vessels. N=3; scale bars = 20uM.

**Figure S2 – Rab11 is deleted in the E14.5 pancreatic epithelium of the Rab11$^{pancDKO}$.** E14.5 pancreas tissue was stained in section for nuclei (DAPI, blue), E-Cadherin (white), Muc1 (magenta) and Rab11A (green). In control tissue, Rab11A is strongly enriched at the apical lumenal membrane marked by Muc1 (A-A''). By contrast, there is a loss of Rab11A signal in the Rab11$^{pancDKO}$ tissue (B-B''). n>3, scale bars =10uM.

**Figure S3 – Loss of Rab11 leads to pancreatic hypoplasia in late gestation and postnatally.** At dissection at both E12.5 and E14.5, there are no obvious defects in pancreatic morphogenesis (A-B shown on stomach; myr-td-Tomato, cyan) (C-D shown on stomach; cyan outline). At E18.5, mutant embryos do not appear grossly different from their control littermates (E), but there is clear pancreatic hypoplasia in Rab11$^{pancDKO}$ embryos (F-G'; cyan outline). Additionally, we observed cystic-like structures, marked by yellow arrowheads (G'). These defects are exacerbated by P2 when comparing a mutant that was already visibly failing to thrive to a control littermate (H-I'). The pancreas is severely hypoplastic (red outlines) and is similarly accompanied by cyst-like structures (yellow arrowheads). n=7 for E12.5; n>>3 for E14.5; n>3 for E18.5; n=3 for postnatal; scale bars A-D=0.1cm, F-G', H, I=0.25cm, H', I'=25uM; ruler in (E) show centimeters.

**Figure S4 – Lumens in Rab11$^{pancDKO}$ pancreata arise from cells that retain Rab11A expression.** At E18.5 in tissue sections, there are instances of complete (B-B'') and partial (C-C'') deletion of Rab11A as compared to controls (A-A'') (DAPI, blue; E-Cad, white; Muc1, magenta; Rab11A, green). In the few open lumens present in the mutants, the ductal structures surrounding them retain expression of Rab11A due to incomplete deletion of Rab11A by Pdx1-Cre. n>3; scale bars=10uM.

**Supplementary Table 1**

Antibodies used in presented studies

| Antigen | Host | Concentration | Manufacturer | Catalog # |
|---|---|---|---|---|
| **Mucin1** | Armenian Hamster | 1:500 | ThermoFisher | HM-1630 |
| **Rab11A** | Rabbit | 1:50 | US Biologicals | R0009-05 |
| **E-Cadherin** | Mouse | 1:500 | BD Trans | 610182 |
| **CPA1** | Goat | 1:100 | R&D | AF2765 |
| **aPKC** | Rabbit | 1:100 | Santa Cruz | sc-216 |
| **Sox9** | Rabbit | 1:100 | Millipore | AB5535 |
| **Insulin** | Rabbit | 1:100 | Cell Signaling | 3014S |
| **Glucagon** | Rabbit | 1:100 | Cell Signaling | 2760S |
| **phospho-Histone H3** | Rabbit | 1:100 | Millipore | 06-570 |
| **Par3** | Rabbit | 1:100 | Millipore | 07-330 |
| **Podocalyxin** | Goat | 1:100 | R&D | AF1556 |
| **GM130** | Rabbit | 1:100 | AbCam | ab52649 |
| **Laminin** | Rabbit | 1:100 | Sigma | L9393 |
| **Par6b** | Rabbit | 1:100 | Santa Cruz | SC-67392 |
| **Crumbs3** | Rabbit | 1:100 | Sigma | HPA013835 |
| **ZO-1** | Mouse | 1:100 | Invitrogen | 33-9100 |
