## Supplemental Figures for "Rab11 is essential to pancreas morphogenesis, lumen formation and endocrine mass"

### Figure S1 – Rab11 is expressed in all pancreatic lineages at E18.5.

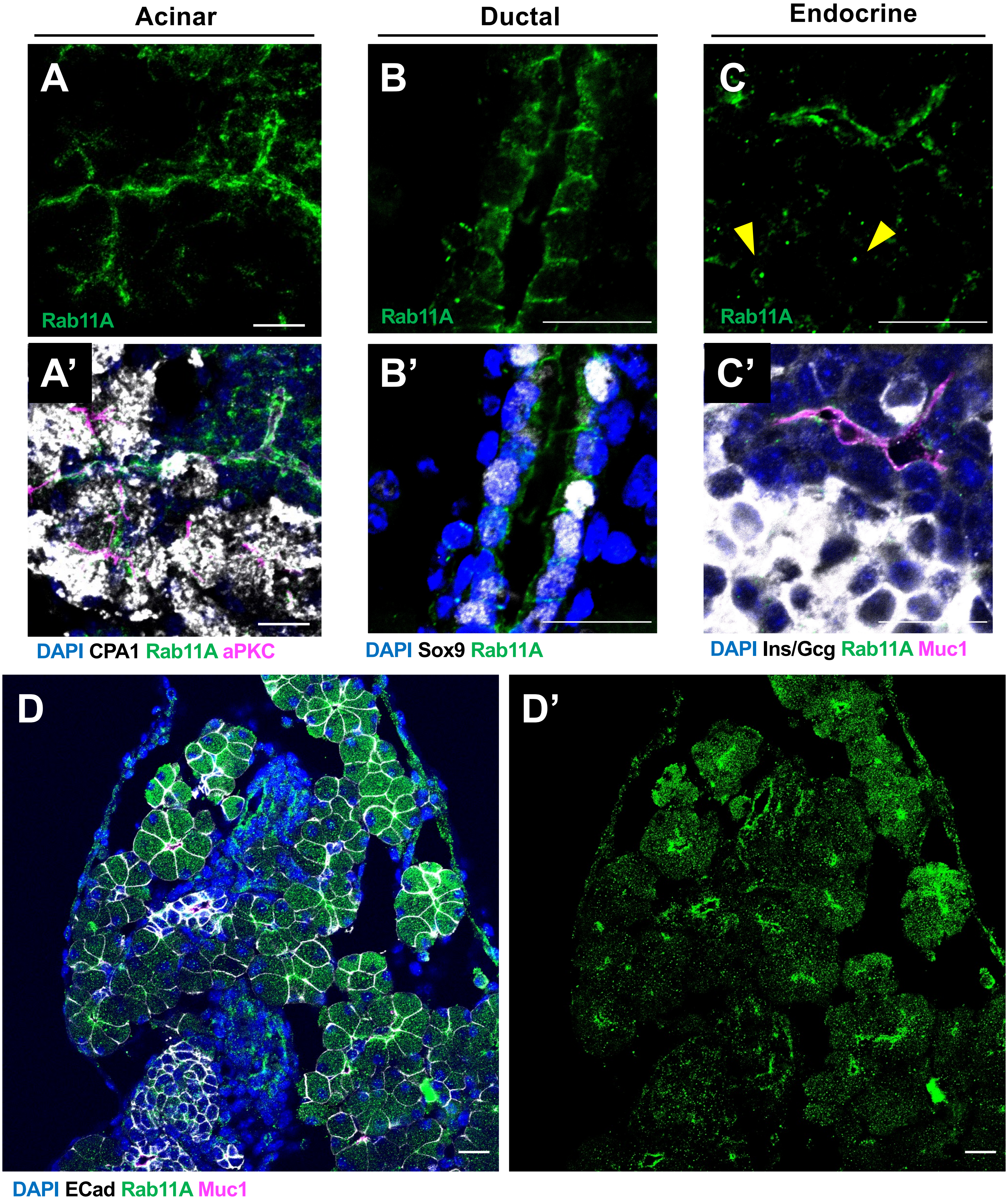

**Figure S2 – Rab11 is deleted in the E14.5 pancreatic epithelium of the Rab11$^{pancDKO}$.**

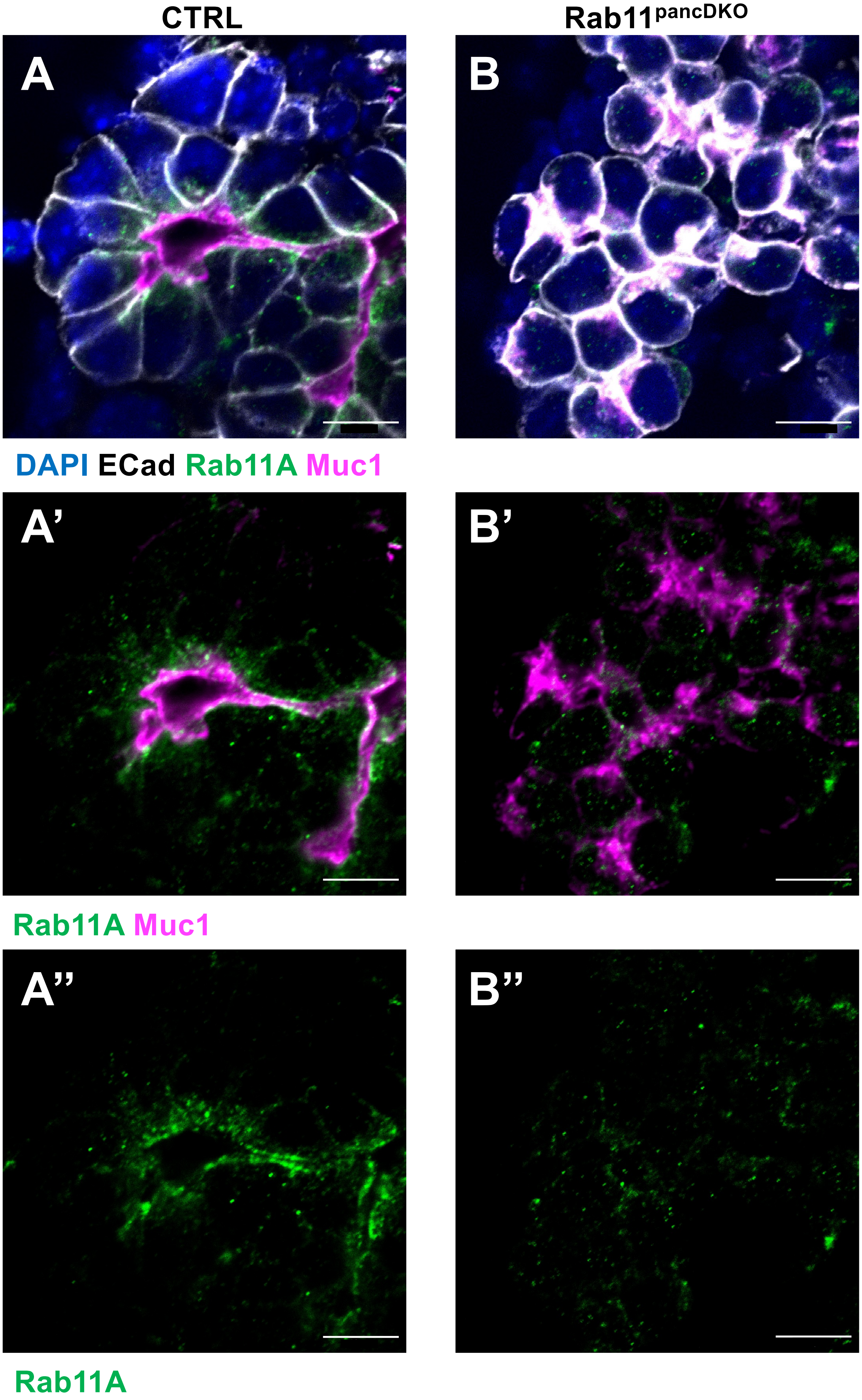

**Figure S3 – Loss of Rab11 leads to pancreatic hypoplasia in late gestation and postnatally**

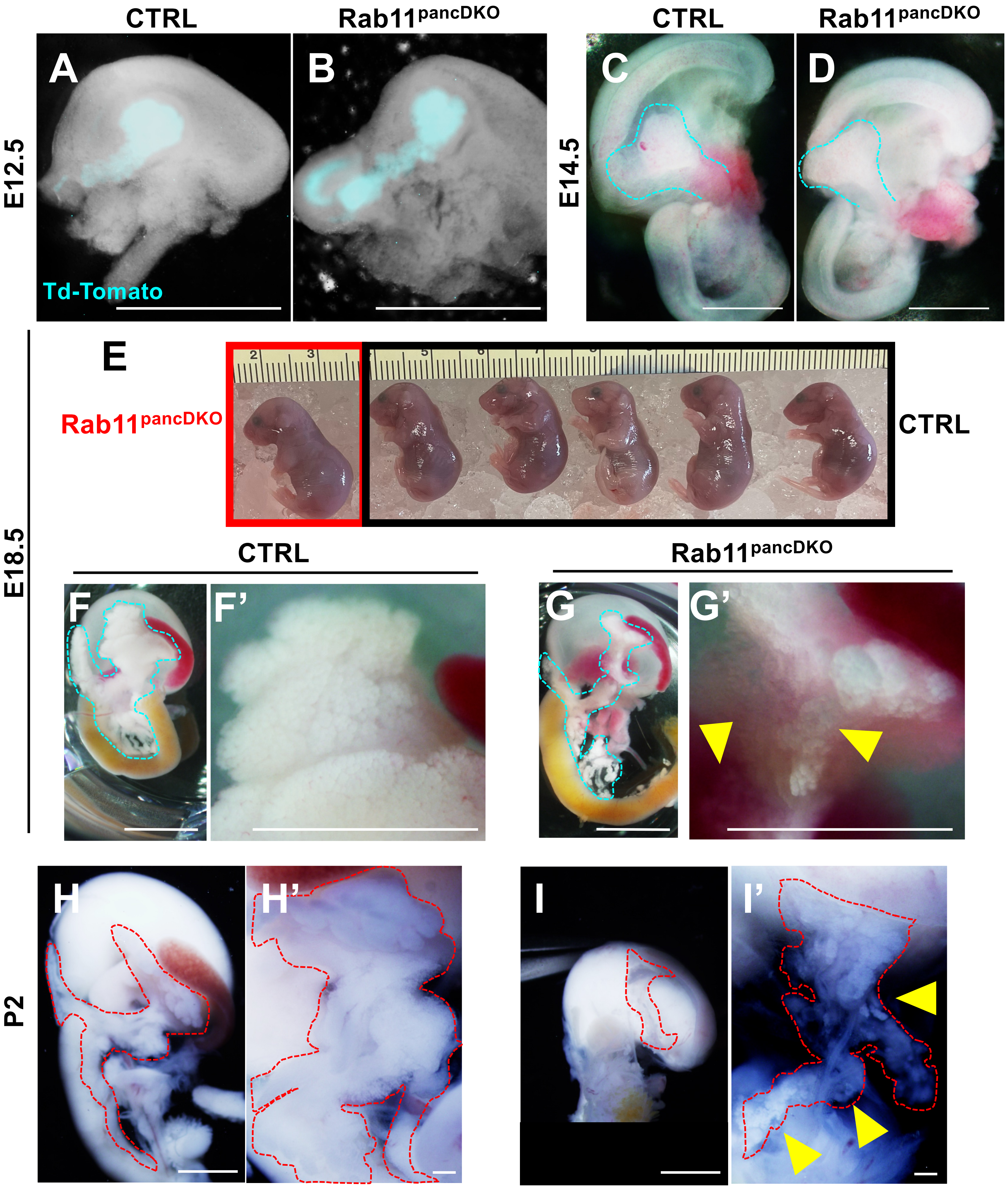

**Figure S4 – Lumens in Rab11$^{pancDKO}$ pancreata arise from cells that retain Rab11A expression.**

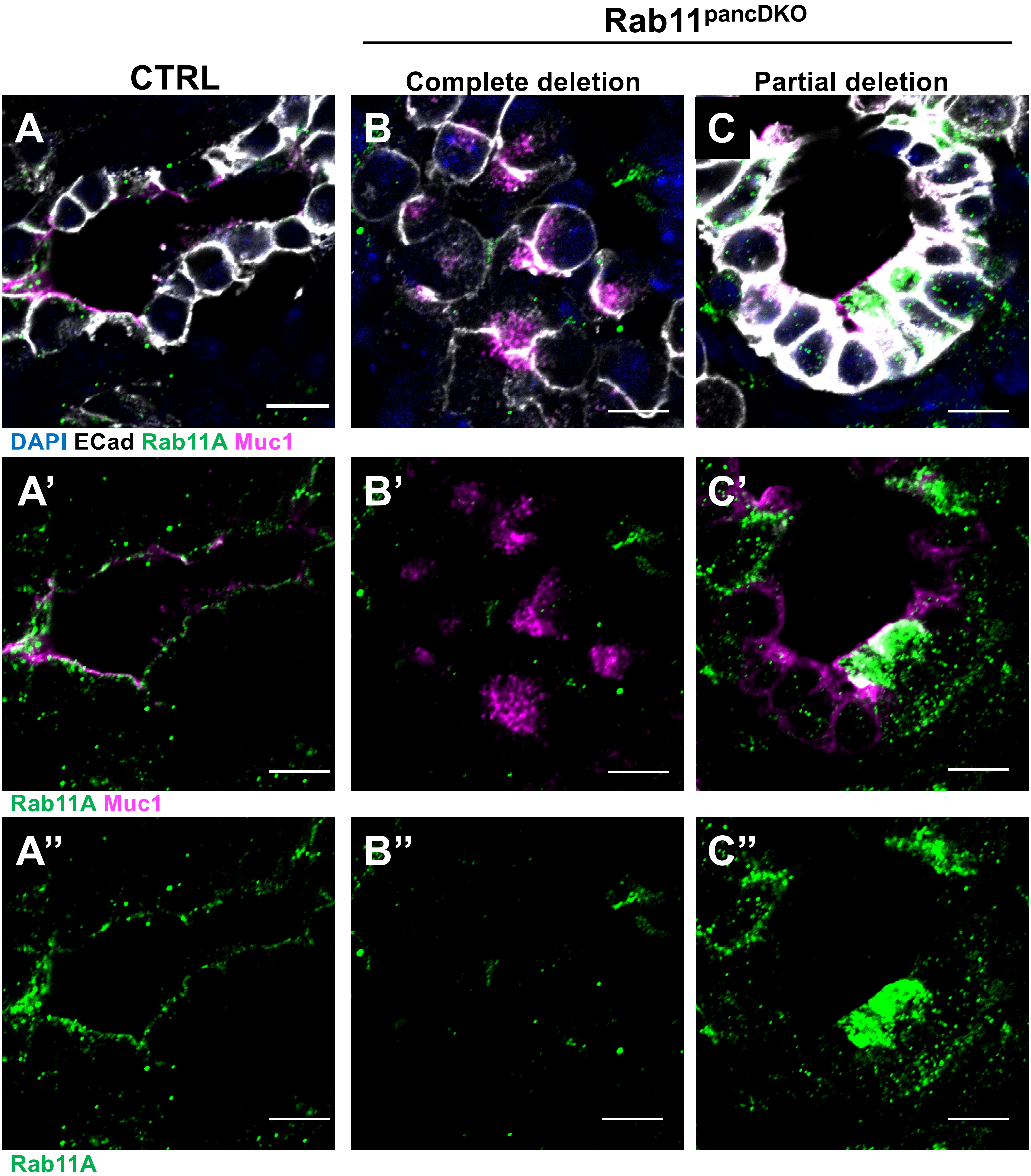
